# The Queensland Twin Adolescent Brain Project, a longitudinal study of adolescent brain development

**DOI:** 10.1101/2022.05.19.492753

**Authors:** Lachlan T. Strike, Narelle K. Hansell, Kai-Hsiang Chuang, Jessica L. Miller, Greig I. de Zubicaray, Paul M. Thompson, Katie L. McMahon, Margaret J. Wright

## Abstract

We describe the Queensland Twin Adolescent Brain (QTAB) dataset and provide a detailed methodology and technical validation to facilitate data usage. The QTAB dataset comprises multimodal neuroimaging, as well as cognitive and mental health data collected in adolescent twins over two sessions (session 1: N = 422, age 9-14 years; session 2: N = 304, 10-16 years). The MRI protocol consisted of T1-weighted (MP2RAGE), T2-weighted, FLAIR, high-resolution TSE, SWI, resting-state fMRI, DWI, and ASL scans. Two fMRI tasks were added in session 2: an emotional conflict task and a passive movie-watching task. Outside of the scanner, we assessed cognitive function using standardised tests. We also obtained self-reports of symptoms for anxiety and depression, perceived stress, sleepiness, pubertal development measures, and risk and protective factors. We additionally collected several biological samples for genomic and metagenomic analysis. The QTAB project was established to promote health-related research in adolescence.

## Background & Summary

Adolescence is critical for understanding brain changes associated with depression^1-3^, as nearly half of lifetime diagnoses begin by age 14^4^. Adolescents who experience depression are more likely as adults to have poor mental and physical health, lower levels of educational attainment, lower salaries, and more relationship difficulties^5-7^. During adolescence, the brain’s cognitive control, emotion, and reward-related circuitries are undergoing substantial development^8,9^ – developmental changes that can be impaired by restricted sleep, which is common among teenagers^10^. In addition, hormonal surges and consequent physical maturation linked to pubertal development in adolescence are thought to affect multiple aspects of brain development, social cognition, and peer relations^11^.

Sex differences in mental health problems emerge in puberty, a developmental period characterised by rapid increases in estrogen in girls and testosterone in boys. Before puberty, depression occurs relatively infrequently in girls and boys. After puberty, there is a sharp increase in the incidence of depression, with adolescent girls around twice as likely to experience depression as boys^12^. Children who start puberty earlier are at greater risk for depressive symptoms and anxiety, especially girls^13^. Adolescent boys, by contrast, are more likely than girls to develop substance use disorders and die by suicide^14^. Importantly, these pubertal hormones cross the blood-brain barrier, influence brain development, and affect various signalling pathways (e.g., neurotransmitter activity) that underlie mood and cognition^15^. Thus, puberty involves transformation across virtually every psychobiological domain—endocrine, neural, physical, cognitive, and socioemotional—and represents a vulnerable time during which depressive symptoms and psychiatric conditions may emerge.

Magnetic resonance imaging (MRI) allows developmental brain changes to be studied *in vivo* while retaining the advantage of being non-invasive, thus providing an invaluable tool for longitudinal research in a population sample. In recent years, terrific progress has been made towards characterizing typical adolescent structural and functional development^1,2,8,16,17^ and understanding brain structure and function changes during adolescence associated with various mental health disorders^1-3,16^. However, there is still little work in this sensitive period of neurodevelopment using genetically informative samples. Further, prior genetic studies have primarily been cross-sectional, and we lack knowledge of how this critical transitional period between childhood and adulthood contributes to optimal cognitive and emotional functioning as well as vulnerability to brain disorders and mental illness. Thus, longitudinal studies are vital to characterise the structural and functional integrity of the brain in genetically informative adolescent samples.

To this end, we present the Queensland Twin Adolescent Brain (QTAB) dataset: a multimodal neuroimaging dataset of Australian adolescent twins with mental health, cognition and social behaviour data collected over two time points (i.e., sessions). Comparisons within and between identical (MZ) and non-identical (DZ) twin pairs -further powered by multiple assessments - provide a rich information source about genetic and environmental contributions to developmental associations and enable stronger tests of causal hypotheses than do comparisons involving unrelated adolescents^18^.

The QTAB project reflected a concerted effort to:

1. Assess brain development in a large population sample of adolescent twins and disentangle the influence of genetic and environmental factors on neurodevelopmental trajectories.
2. Investigate whether neurodevelopmental trajectories are the same for adolescent boys and girls and whether any differences are linked with pubertal maturation and genetic and environmental factors.
3. Concurrently assess cognitive function to examine how neurodevelopmental changes are associated with the acquisition of complex tasks (e.g., working memory) and behaviour (e.g., emotional functioning) and whether shared genetic or environmental factors underpin associations.
4. Concurrently assess psychiatric symptoms and examine whether those with anxiety or depressive symptoms follow a different developmental pathway, whether they exhibit early or delayed brain maturation, and whether genetic and environmental factors influence these atypical developmental trajectories.

Here, we make this multimodal QTAB dataset publicly available and provide an overview of the collected imaging, mental health, cognition, and social behaviour data. In doing so, we hope to make a significant and novel contribution to our understanding of the numerous emotional and behavioural health problems that emerge during the developmental period of adolescence. At the same time, we believe the QTAB dataset provides a significant opportunity for combining data with existing adolescent studies such as the Adolescent Brain Cognitive Development (ABCD) ^17^, Lifespan Human Connectome Project in Development (HCP-D) ^19^, and the NKI-Rockland Sample Longitudinal Discovery of Brain Development Trajectories^20^ projects, as well as consortia efforts such as ENIGMA Lifespan^21,22^.

## Methods

### Recruitment of the QTAB Cohort

Participants were recruited through local and national twin registries (i.e., The Queensland Twin (QTwin) Register [https://www.qimrberghofer.edu.au/qtwin] and Twin Research Australia [TRA] [https://www.twins.org.au]) or via flyers and online postings (study website). Families were invited to participate if the twins were aged 9 to 13 years (i.e., year of birth 2004-2010) and lived within approximately 2 hours travelling distance of the study centre in Brisbane (i.e., South-East Queensland). Four hundred and forty families met the inclusion criteria (QTwin Register: 356 families; TRA: 48; study website: 36), of which 63 (14% of families) declined on the first approach (letter followed by phone call) and a further 166 families were excluded (16% excluded on screening; 22% passive refusals). Exclusion criteria included serious medical, neurological, psychiatric (including diagnosis of autism spectrum disorder [ASD] or attention deficit hyperactivity disorder [ADHD]), or cardiovascular conditions, history of a serious head injury, and any cognitive, physical, or sensory challenges that would limit ability to understand or complete procedures. In addition, both twins (i.e., co-twins) had to have no MRI contraindications (e.g., metal implants or artefact inducing orthodontic braces). Thus, the total available sample comprised 211 families, including 206 twin pairs and 5 triplet sets. Only two individuals from each trio were scanned, and hereafter are referred to as a twin pair.

### QTAB Protocol and Participants

Baseline measures (i.e., session 1 data) were collected between June 2017 and October 2019, followed by a second wave of data collection (i.e., session 2 data) from November 2019 to January 2021. Both sessions, each taking ≈3.5 hours, included structural and functional brain imaging, assessments of pubertal status, cognition, anxiety and depressive symptoms plus risk and protective factors, sleep-wake behaviour, and the collection of several biological samples (Fig. 1). While the target interval between sessions was ≈18 months, to enable as many participants as possible to return for their second assessment, and because of the interruption to data collection due to the COVID-19 pandemic, we allowed for some latency jitter, which resulted in an interval of 13 to 30 months (mean 20 months).

**Fig. 1.**
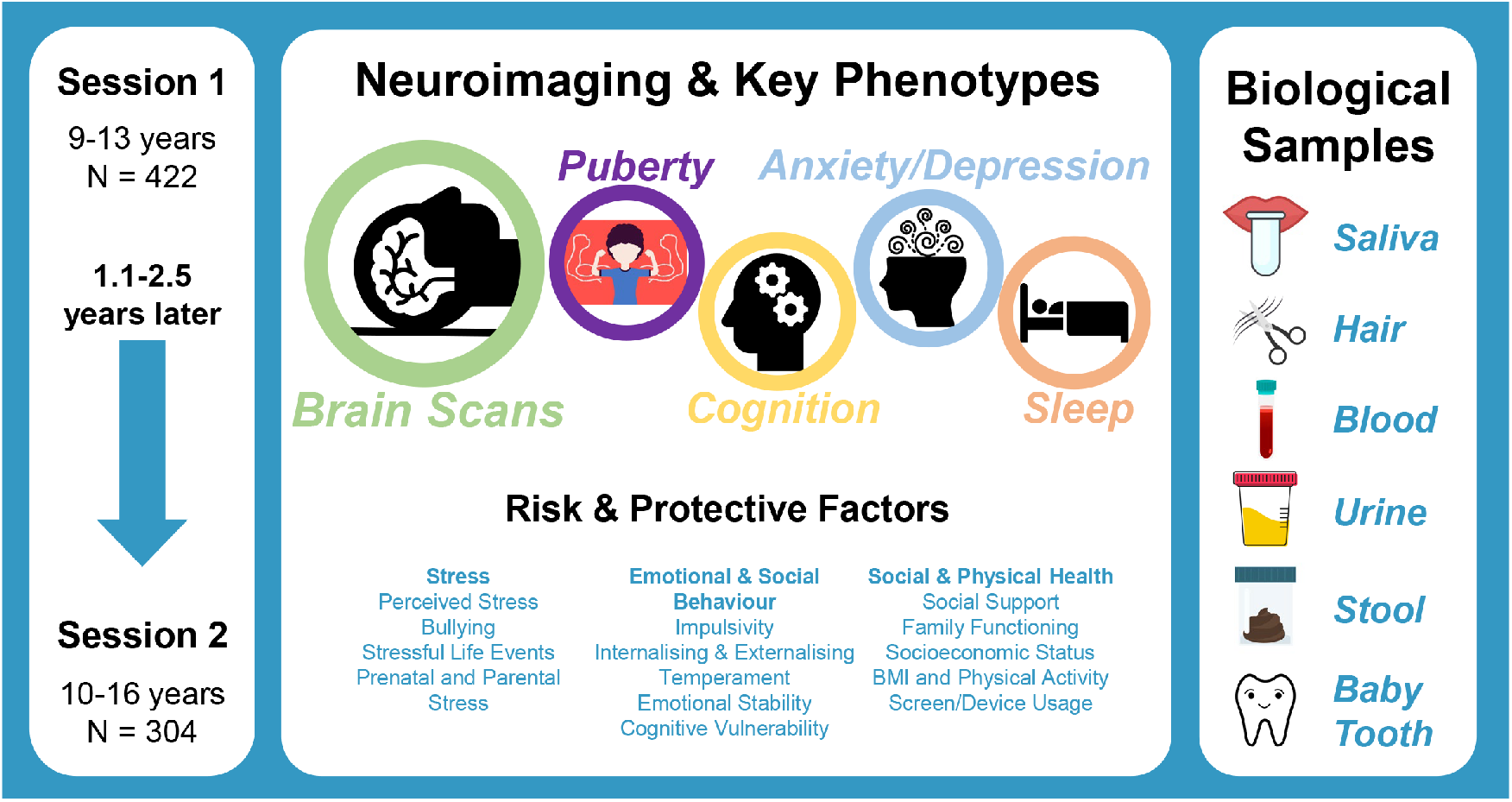
Data collection overview. Data from imaging and questionnaire/cognition components were collected concurrently between twin pairs (i.e., while one twin underwent brain imaging, their co-twin completed the mental health/behavioural questionnaire and cognitive testing). An early morning saliva sample was collected at home and brought to the session, along with any available baby teeth. Hair, blood, and urine samples were obtained during the session, with faecal samples and accelerometry measures of sleep behaviour collected at home. Neuroimaging and key phenotypes, saliva and hair samples were collected at sessions 1 and 2.

Co-twins attended the study centre together and were assessed in parallel, with one twin undergoing brain imaging while the other completed assessments outside of the scanner. In addition, the parent accompanying the twins - usually the mother (94% of families) - completed a parent questionnaire providing family information and parental assessments of the twins. Hair, blood, and urine samples were obtained during the session. Early morning saliva samples were collected at home on the morning of their session visit and brought to the study centre, as were any available baby teeth. At the end of the session, the twins were provided with an accelerometer to objectively measure their sleep behaviour for two weeks and a stool collection pack. The accelerometer and stool sample were returned to the study centre by post and courier, respectively. Informed written consent, including data sharing permission, was obtained from both the twins and the attending parent; families were compensated $150 (AUD) per session. Ethics approval for this study was obtained through the Children’s Health Queensland Human Research Ethics Committee (HREC) (reference HREC/16/QRCH/270) and the University of Queensland HREC (reference 2016001784).

Five participants were unable to be scanned in session 1 due to a reluctance to enter the MRI system. Thus, the imaging dataset at session 1 consists of a maximum of 417 participants (48% female, mean age 11.3 ± 1.4 years, age range 9.0 to 14.4 years, 82% right-handed), including 206 twin pairs (109 MZ, 97 DZ) and 5 unpaired twins. In session 2, we focused on pairs with good quality images at session 1 (i.e., both twins had good T1w scans). In total, 59 families did not return for session 2 (39 were no longer eligible (i.e., braces, moved interstate and/or poor T1w scans at session 1) and 20 declined), so the dataset was limited to 304 participants (52% female, mean age 13.04 ± 1.52 years, age range 10.1 to 16.4 years, 85% right-handed), comprising of 152 twin pairs (81 MZ and 71 DZ). Key demographics for QTAB participants are presented in Table 1.

**Table 1.** Key demographics for QTAB participants (including family and birth information). Notes: **Handedness** was based on participant self-report. **Zygosity** in same-sex twins was determined by genotypic data (93% of twin pairs) or parental questionnaire (7% of twin pairs). Genome-wide identity-by-descent (IBD) was estimated for twin pairs using PLINK^23^, after quality checking procedures^24^. Pairs whose proportion IBD (PI_HAT) was greater than 0.99 were classified as MZ pairs; remaining pairs (whose proportion IBD [PI_HAT] was between 0.41 and 0.61) were classified as DZ pairs. **School grade year** was classified as follows: Year 3-4 aged 8–10yrs (primary school), Year 5-6 aged 10–12 (last 2yrs of primary school), Year 7-9 aged 12–15 (middle/high school), Year 10-11 aged 15-17 (high school). **Ancestry** was based on self-perceived group/cultural identification with ancestry options obtained from the 2016 Australian Census. Each family identified 1 to 7 cultural groups (M=2.1±0.9). The most-reported ancestries (as a proportion of all reported ancestries) were Australian (30.6%) and English (24.2%), with the most reported non-Caucasian ancestry being Chinese (1.3% of all reported ancestries). Ancestries reported for the Australian population in the 2016 census were similar (33.5% Australian, 36.1% English, and 5.6% Chinese). **Socioeconomic status** refers to Socio-Economic Index for Areas (SEIFA) Index of Relative Socio-Economic Advantage and Disadvantage (IRSAD) deciles (low = deciles 1-3, average = deciles 4-7, high = deciles 8-10). **Education** (parent self/partner report) classifications are based McMillan et al. ^25^, with post-school diploma and trade Certificate categories collapsed into “Trade”. **Birth information** was provided through parental report. The birth statistics available for QTAB (note that birth weight is not available for over half the sample) are similar to those for twins born in Australia between 2001 and 2010^26^, of whom 53.6% were born preterm, while 50.2% were low birthweight, and 8.7% were very low birthweight. QTAB maternal age was also similar to national statistics (reported as 2.0/33.2/60.2/4.7%^26^, based on the age categories in Table 1). An Assisted Reproductive Technology (ART) reported rate of 19.9% for QTAB twins born 2004-2010 is consistent with the 23.3% reported for 9,831 twin deliveries in Australia from 2007 to 2011^27^. Abbreviations: *DZ* dizygotic; *IVF* in vitro fertilisation; *MZ* monozygotic; *NA* not available; *Univ* university.

| Participants | Session 1 | Session 2 |
| --- | --- | --- |
| **N** | 422 | 304 |
| **Age** (years) | 11.3±1.4 | 13.0±1.5 |
| **Sex** (% biological female/male) | 48.3/51.2 | 51.6/48.4 |
| **Handedness** (% right/left) | 82.2/17.8 | 84.9/15.1 |
| **Zygosity** (% MZ/DZ) | 52.6/47.4 | 53.3/46.7 |
| **School Grade Year** (% Y3-4/Y5-6/Y7-9/Y10-11) | 22.3/46/31.7/0 | 2.6/31.6/56.9/8.9 |
| Families |  |  |
| **N** | 211 | 152 |
| **Ancestry** (group identification as % of all reported groups) |  |  |
| Most Common: Australian/English | 30.6/24.1 | 28.8/24.1 |
| Non-Caucasian: All/Asian/First Nations Australian | 6.9/3.4/0.9 | 5.6/3.1/0.9 |
| **Socioeconomic Status** (% low/average/high) | 9.0/34.1/56.9 | 9.2/35.5/55.3 |
| **Education** |  |  |
| Mother (% Univ/Trade/School/NA) | 52.6/28.9/18.5/0 | 51.3/27.7/21.1/0 |
| Father (% Univ/Trade/School/NA) | 39.3/32.7/22.2/5.8 | 40.7/31.5/22.4/5.4 |
| **Birth Information** |  |  |
| Gestational age (% preterm (<37 wks)/NA) | 54.5/1.9 | 52.6/1.3 |
| Weight (% low: 1500-2499g/very low: <1500g/NA) | 36.5/5.7/57.8 | 36.5/5.6/57.9 |
| Maternal age in years (<20/20-29/30-39/≥40/NA) | 0.9/30.3/62.6/3.3/2.8 | 1.3/29.6/62.5/3.9/2.6 |
| % Assisted Reproduction Technology (e.g., IVF) | 19.9 | 19.7 |

### Image Acquisition

Structural and functional scans were acquired on a 3T Magnetom Prisma (Siemens Medical Solutions, Erlangen) paired with a 64-channel head coil at the Centre for Advanced Imaging, University of Queensland. Session 1 scans were collected in a fixed order: localisers, 3D T1-weighted, two runs of resting-state fMRI (rs-fMRI), four runs of diffusion-weighted imaging (DWI), three runs of high-resolution Turbo Spin Echo (TSE) imaging, 3D fluid-attenuated inversion recovery (FLAIR), 3D T2-weighted, Arterial Spin Labelling (ASL), and 3D susceptibility-weighted-imaging (SWI). In session 2, scan order was very similar, with the addition of two task-based fMRI scans midway through the imaging session, i.e., localisers, T1-weighted, two runs of rs-fMRI, four runs of DWI, two runs of TSE, passive movie-watching task (Partly Cloudy), emotional conflict fMRI task, ASL, SWI, FLAIR, T2w. Immediately after each scan, the images were checked for apparent artifacts (e.g., head motion), and scans were re-acquired if necessary. Scan parameters are summarised in Table 2. Protocol optimisations (e.g., increase in the number of TSE volumes to improve hippocampus and amygdala coverage) and sequence reissues (e.g., to the T1w) that occurred across participants or sessions are noted as a protocol code (detailed in Table 2 and provided in *participants*.*tsv*; see *Data Records*). MRI acquisition took approximately 1 hour in session 1 and 1:15 hours for session 2.

**Table 2.** MRI acquisition parameters for the QTAB project. Notes: Voxel size was optimised for each modality. The scanning protocol code for each participant is defined in *participants*.*tsv* (see *Data Records*). Abbreviations: *A>>P* anterior-posterior phase encoding; *deg* degree; *FA* flip angle; *FOV* field of view; *ms* milliseconds; P*>>A* posterior-anterior phase encoding; *TE* time to echo; *TI* inversion time; *TR* repetition time <index> refers to run number. * Full NifTI filename begins with sub-<participant_id>_ses-<session_id>

| Modality | Voxel Size | Key Parameters | NIfTI Filename |
| --- | --- | --- | --- |
| **Anatomical** |  |  |  |
| T1w | 0.8 x 0.8 x 0.8 mm | 3D MP2RAGE, sagittal acquisition, TR/TE = 4000/2.99 ms, T1/TI2 = 700/2220 ms, FA1/FA2 = 6/7 deg, slices = 192, FOV = 256 x 240 mm, in-plane acceleration = 3, duration = 6:04 min:sec. *Protocol change:* T1w reissue during session 2 (protocol code 2c). | *_inv-1_MP2RAGE,<br>*_inv-2_MP2RAGE,<br>*_UNIT1,<br>*_UNIT1_denoised |
| T1w (reissue) |  | As above, except: TE = 3.03 ms |  |
| T2w | 1.0 x 1.0 x 1.0 mm | 3D T2, sagittal acquisition, TR/TE = 3200/408 ms, slices = 176, FOV = 256 x 256 mm, in-plane acceleration = 2, duration = 3:31 min:sec | *_T2w |
| FLAIR | 1.0 x 1.0 x 1.0 mm | 3D T2-FLAIR, sagittal acquisition, TR/TE = 5000/388 ms, slices = 160, FOV = 256 x 256 mm, in-plane acceleration = 2, duration = 5:55 min:sec | *_FLAIR |
| TSE | 0.5 x 0.5 x 1.0 mm (interpolated in k-space to 0.25 x 0.25 x 1.0 mm) | 2D TSE slab orthogonal to the hippocampus, TR/TE = 8460/67 ms, FA = 150 deg, slices = 60, FOV = 192 x 192 mm, duration = 4:48 min:sec (each). *Protocol change:* Number of slices was initially 50, increased to 55 to improve hippocampal coverage (protocol code 1b), increased again to 60 to improve amygdala coverage (protocol code 1c). | *_T2w_TSE_run-<index> |
| **Susceptibility** |  |  |  |
| SWI | 1.1 x 1.1 x 2.0 mm | 3D SWI, TR/TE = 27/20 ms, FA = 15 deg, slices = 60, FOV = 220 x 198 mm, in-plane acceleration = 2, duration = 2:51 min:sec | *_swi, *_part-mag_GRE,<br>*_part-phase_GRE, *_minIP |
| **Diffusion** |  |  |  |
| dMRI | 2.0 x 2.0 x 2.0 mm | 4 acquisitions (2 A>>P, 2 P>>A), TR/TE = 3800/70 ms, FA = 90 deg, in-plane acceleration = 2, slice acceleration = 2, slices = 68, FOV = 244 x 244 mm, b-values: 0 (x 3), 1000 (x 5), 3000 (x 15) s/mm<sup>2</sup>, duration = 1:47 min:sec (each) | *_dir-AP_run-<index>_dwi,<br>*_dir-PA_run-<index>_dwi |
| **Functional** |  |  |  |
| rs-fMRI | 2.0 x 2.0 x 2.0 mm | 2 acquisitions (1 A>>P, 1 P>>A), TR/TE = 930/30 ms, FA = 52 deg, slices = 72, FOV = 208 x 208 mm, slice acceleration = 6, volumes = 327 (each), duration = 5:13 min:sec (each) | *_task-rest_dir-AP_bold,<br>*_task-rest_dir-PA_bold |
| fMRI | 2.4 x 2.4 x 2.4 mm | TR/TE = 800/30 ms, FA = 52 deg, A>>P, slices = 60, FOV = 216 x 216 mm, slice acceleration = 6, volumes = 380 (Partly Cloudy), volumes = 837 (emotional conflict), duration = 5:11 min:sec (Partly Cloudy), duration = 11:17 min:sec (emotional conflict). *Protocol change:* number of volumes in the emotional conflict task was initially 830, increased to 837 (protocol code 2b). | *_task-partlycloudy_bold,<br>*_task-emotionalconflict_bold |
| **Perfusion** |  |  |  |
| ASL | 1.7 x 1.7 x 4.0 mm | 2D PCASL, TR/TE = 4800/13 ms, slices = 24, labelling time = 1500ms, post-labelling delay = 1500ms, volumes = 31 (1 x M0, 15 label-control pairs), slices = 24, FOV = 220 x 220 mm, duration = 2:30 min:sec | *_asl |
| ASL M0 | 1.7 x 1.7 x 4.0 mm | As above, except TR = 8000 ms, volumes = 3, duration = 0:26 min:sec (session 1), 0:34 min:sec (session 2) | *_m0scan |
| **Field Maps** |  |  |  |
| Field maps | 2.4 x 2.4 x 2.4 mm | 2 acquisitions (1 A>>P, 1 P>>A), TR/TE = 6350/63 ms, slices = 60, FOV = 216 x 216 mm, volumes = 4 (each), duration = 0:33 min:sec (each) | *_dir-AP_epi,<br>*_dir-PA_epi |

### Image Modalities

#### Anatomical

A 3D whole-brain T1-weighted image was acquired using a work-in-progress Magnetisation Prepared with 2 Rapid Gradient Echoes (MP2RAGE) sequence. The MP2RAGE sequence combines two individual images acquired at different inversion times (inv-1_MP2RAGE, inv-2_MP2RAGE) to produce an image corrected for B_1_ and T2* inhomogeneities^28,29^. This “uniform” image (UNIT1) improves T_1_ contrast compared to conventional T1-weighted protocols. However, a by-product of the MP2RAGE protocol is an amplification of background noise (also known as “salt and pepper” noise), which is problematic for automatic registration and segmentation algorithms (see *Usage Notes* for techniques to address this). The sequence generates a denoised uniform image (UNIT1_denoised), though some background noise is still present in this image. The MP2RAGE sequence was reissued during session 2, producing a denoised uniform image (UNIT1_denoised) without background noise.

T2-weighted and FLAIR whole-brain 3D sequences were acquired to examine brain pathology and to improve the visibility of brain regions bordering CSF (e.g., cortical pial surface, periventricular structures). Due to time constraints, the T2-weighted and FLAIR sequences were excluded for some participants (≈5% in session 1, ≈10% in session 2). T2-weighted TSE scans (slab aligned orthogonally to the hippocampus) were acquired to examine the structural properties of the hippocampus and amygdala at the subdivision level (three runs collected at session 1, reduced to two runs at session 2).

#### Susceptibility

SWI scans were acquired to study the pathological mechanisms underlying paediatric brain diseases and disorders. In addition to a combined magnitude and phase image (*_swi), the SWI sequence produces magnitude (*_part-mag_GRE), phase (*_part-phase_GRE), and minimum intensity projection (*_minIP) images.

#### Diffusion

Multi-shell diffusion weighted images were acquired in four runs with opposing phase encode directions (two A>>P and two P>>A runs). Each run consisted of 23 diffusion weighted volumes (which includes b = 0 s/mm^2^ (3 volumes), b = 1000 s/mm^2^ (5 volumes), b = 3000 s/mm^2^ (15 volumes).

#### Functional

Resting-state scans were acquired consecutively over two runs with reversed-phase encoding directions (anterior-posterior (A>>P), posterior-anterior (P>>A)). Participants were instructed to relax with their eyes open, with an abstract visual stimulus displayed to improve resting-state compliance^30^. The stimulus for functional scans was presented on a back-projection screen that the participants viewed via a mirror attached to the head coil.

In session 2, the emotional conflict task was based on a previously characterised emotional conflict task^31,32^ and probed emotion-relevant neural processes engaging the amygdala and other structures that represent the processing of emotions and faces. In each trial, participants were presented with a face displaying a fearful or happy expression and the words “happy” or “fear” written across the face to create congruent and incongruent emotional stimuli. Participants were asked to identify the facial emotion while ignoring the word overlayed on the face. Responses were made via a button press (left button = fearful face, right button = happy face). Stimuli were presented for 1,000 ms, with a varying interstimulus interval of 2,000 – 4,000 ms (average 3,000 ms; 163 trials in total), presented in a fixed order, counterbalanced across trial types for expression, word, and sex. Participants completed an in-scanner practice task (8 trials) immediately prior to the task. Stimuli were presented using the Cogent toolbox (www.vislab.ucl.ac.uk/cogent_2000.php) implemented in the MATLAB (www.mathworks.com) programming environment.

Session 2 participants also completed a passive movie-watching task involving the short animated film Partly Cloudy^33^ (silent version). This task was selected based on prior work examining cortical and cognitive changes in adolescents’ social development^34,35^. Participants were instructed to remain still and watch the film. The stimulus was transcoded from a PAL format DVD (see *Usage Notes*) and presented using E-Prime (Psychology Software Tools, Pittsburgh, PA); there was no rest period prior to stimulus presentation. We provide condition timings based on a prior study^34^ (see *Data Records* and *Usage Notes*); however, we note that such a naturalistic movie-watching task can be used to examine a range of different cognitive, emotional, and perceptual processes.

#### Perfusion

ASL scans provide a measure of cerebral blood flow without the requirement of contrast agents or radiation. In session 1 we used a work-in-progress 2D pseudo-continuous ASL (PCASL) sequence. The product sequence was available at the start of data collection for session 2. In both sessions, a separate M0 scan was acquired using a shortened version of the ASL sequence to produce an improved measure of M0 for calibration purposes.

### QTAB Non-Imaging Phenotypes

Table 3 provides a detailed list of the scales and tests used to assess *puberty, cognition, anxiety and depressive symptoms, emotional and social behaviours, social support and family functioning, stress, sleep and physical health, early life and family demographic factors, dietary behaviour, and COVID-19 pandemic specific assessments*. In general, the same measures were collected in sessions 1 and 2, and for several scales, we included both adolescent and parent versions. Selection of tests and self-report scales for phenotypic characterisation was based on validation for use with adolescents (i.e., gold standard tasks with known reliability and applicability to the age range of the QTAB cohort) and ease of administration (i.e., suitability or adaptation for online assessment with an iPad or computer, and whether scales and tests were freely available or provided at a modest cost [e.g., NIH Toolbox Cognition Battery^36^]). Tests and scales that were widely used were also prioritised and highly considered.

**Table 3.**
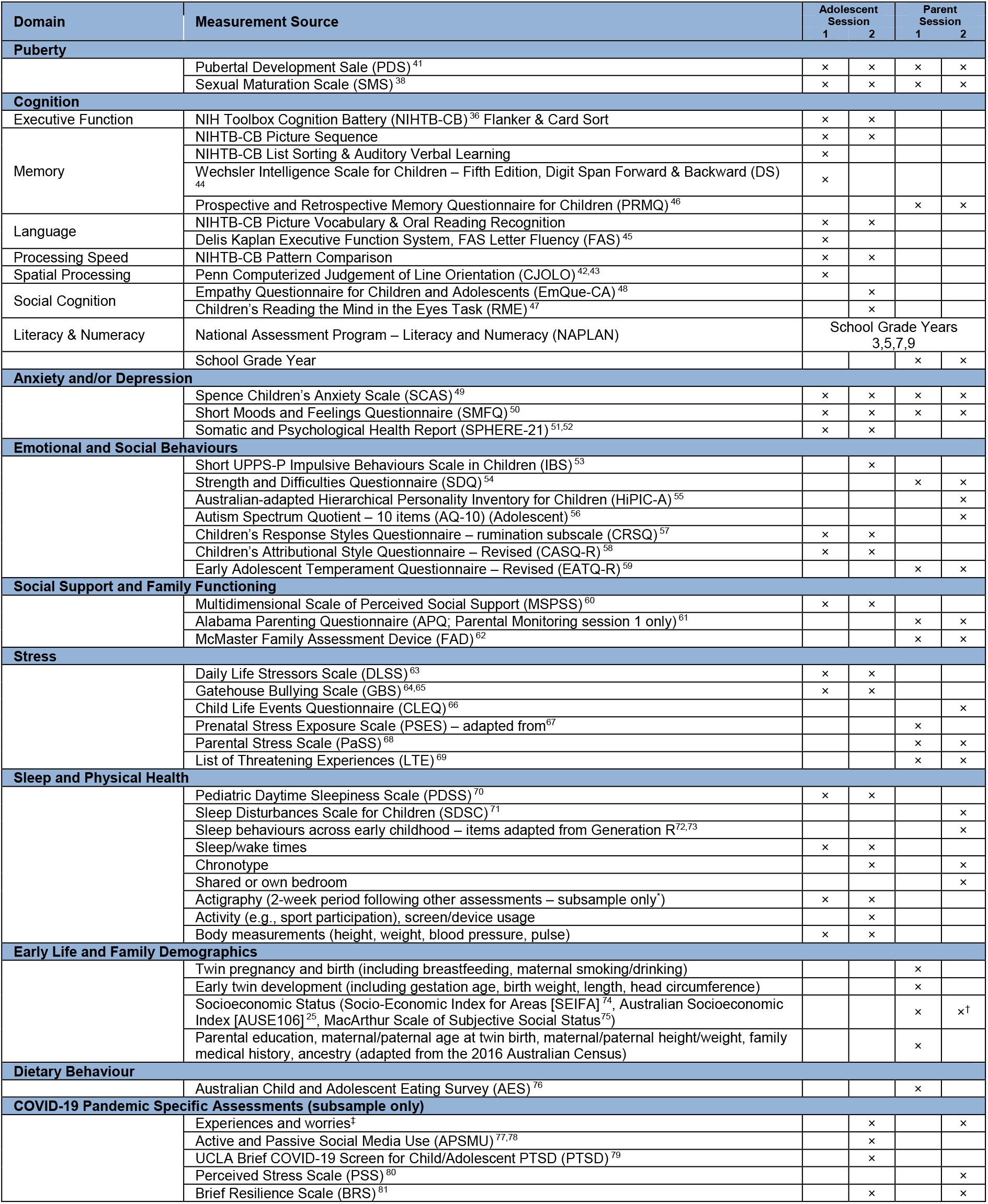
QTAB Non-Imaging Phenotypes (domain and measurement source). ^*^The actigraphy sample was largely restricted (*N* session 1 = 188, session 2 = 134) due to the availability of actigraphy watches rather than a refusal to participate. ^†^SEIFA only. ^‡^ Adapted from multiple COVID-19 survey sources, including the Swinburne University of Technology (Melbourne) and the NIH Office of Behavioral and Social Sciences Research (OBSSR).

#### Puberty

While clinician assessment of sexual maturation is considered superior to self-report, the validation of self-assessment methods has supported their use as an acceptable alternative for research purposes^37^. In QTAB, we assessed pubertal status using a combination of self- and parent-report based on line drawings corresponding to the Tanner stages of pubertal development^38-40^ and complementary questions regarding the emergence of secondary sexual characteristics^41^. Further, to aid in tracking pubertal development, saliva samples were collected at each visit and stored for future analysis of sex hormones (see *Biological Samples* for more information on saliva collection).

#### Cognition

The cognitive tests focussed on 5 domains: Executive Function, Memory, Language, Processing Speed, and Spatial Processing – abilities that are related to general adolescent development, as well as anxiety and depression. For ease of administration, we selected the iPad version of the NIH Toolbox Cognition Battery (NIHTB-CB) ^36^ and the Penn Computerized Judgement of Line Orientation (CJOLO) ^42,43^. Both are now widely used and facilitate the comparison of QTAB data to independent adolescent cohorts, such as the ABCD and HCP-D. The electronic tests were supplemented with 3 well-validated pencil and paper tests from the Wechsler Intelligence Scale for Children – Fifth Edition^44^ (Digit Span Forward & Backward) and Delis Kaplan Executive Function System^45^ (FAS Letter Fluency), as well as the parental report Prospective and Retrospective Memory Questionnaire for Children (PRMQ) ^46^. In session 2, a novel inclusion was the assessment of social cognition using the Children’s Reading Mind in the Eyes Task (RME) ^47^ and the Empathy Questionnaire for Children and Adolescents (EmQue-CA) ^48^. In addition, we obtained consent to access their National Assessment Program – Literacy and Numeracy (NAPLAN) scores (https://www.nap.edu.au/home). NAPLAN is a standardised national assessment of reading, spelling, grammar and punctuation, writing, and numeracy skills undertaken by Australian children in grades 3, 5, 7, and 9 in both government and non-government schools. Currently, NAPLAN scores are not provided as part of the QTAB dataset; please contact authors Greig de Zubicaray and Katie McMahon.

#### Mental Health

We focused on symptoms of *anxiety* and *depression*, both of which increase in prevalence during adolescence. *Depressive* symptoms were assessed by self-report with the Short Moods and Feelings Questionnaire (SMFQ) ^50^ and Somatic and Psychological Health Report (SPHERE-21) ^52^. *Anxiety* symptoms were measured using the Spence Children’s Anxiety Scale (SCAS) ^49^. We also assessed traits linked to mental health, which may indicate risk of progression or vulnerability. *Emotional and social behaviour* measures included: *impulsivity* (Short UPPS-P Impulsive Behaviours Scale in Children [IBS] ^53^), *internalising and externalising* (e.g., emotional and conduct problem scales from the Strength and Difficulties Questionnaire [SDQ] ^54^), *temperament* (Early Adolescent Temperament Questionnaire-Revised [EATQ] ^59^), *emotional stability* (Australian-adapted Hierarchical Personality Inventory for Children [HiPIC-A] ^55^; also extraversion, amenability, conscientiousness, and imagination), symptoms of *autism* (Autism Spectrum Quotient – 10 items [AQ-10] [Adolescent]) ^56^), and *cognitive vulnerability* (rumination subscale from the Children’s Response Styles Questionnaire [CRSQ] ^57^ and the Children’s Attributional Style Questionnaire-Revised [CASQ-R] ^58^, both of which can improve screening of adolescent depression^82^). These dimensional measures investigate constructs relevant to general adolescent maladaptive behaviours (e.g., problem behaviours, risky and impulsive decision making). In addition, *Social Support and Family Functioning* was assessed with the Multidimensional Scale of Perceived Social Support (MSPSS) ^60^, which captures support from family, friends, and significant other^83^ and two scales answered by the parent – the Alabama Parenting Questionnaire (APQ) ^61^ to capture parenting style and the McMaster Family Assessment Device (FAD) ^62^, which provides both dimensional and overall measures of family functioning^84^. Peer influences and family factors are posited to influence adolescent brain development^85-87^ and moderate adolescent risk of experiencing mental health problems^88-91^.

#### Stress

We included structured assessments to capture *perceived stress* (Daily Life Stress Scale [DLSS] ^63^) and *bullying* (Gatehouse Bullying Scale [GBS] ^64,65^, capturing both overt and covert types of victimisation), supplemented with parental report of *stressful life events* (Child Life Events Questionnaire [CLEQ] ^66^ and List of Threatening Experiences [LTE] ^69^, i.e., stressful life events over the last year) and measures of *prenatal* (Prenatal Stress Exposure Scale [PSES] adapted from^67^) and *parental* (Parental Stress Scale [PaSS] ^68^) stress. Adolescence is a period of increased vulnerability to stressors and to stress-related psychopathology, including anxiety and depression – relationships that may be mediated through the effects of stress on the developing adolescent brain^3,92^.

#### Sleep and Physical Health

During each session, we assessed *sleepiness* using the Pediatric Daytime Sleepiness Scale (PDSS) ^70^, which has robust psychometric properties in adolescents aged 11 to 15 years and is associated with academic achievement and mood. Participants completed a sleep diary, recording their sleep and wake times (i.e., sleep duration) on weekdays and weekends. In addition, we obtained several anthropometric measures (e.g., height, weight). In session 2, we also asked whether they were a morning or evening person (i.e., to provide a proxy measure of chronotype) and assessed general physical activity and device usage (questions from the literature^93-95^ and the Youth Risk Behavior Survey, United States, 2019). Further, the parent provided information about sleep disturbances (Sleep Disturbances Scale for Children [SDSC] ^71^ and sleep behaviours across early childhood items, as assessed in the Generation R study^72,73^).

In addition, at the end of each session, a sub-sample of participants were given a wrist-mounted accelerometry recording device (GENEActiv, Activinsights, Kimbolton, UK) to wear for two weeks on the wrist of their non-dominant hand. Accelerometry devices detect motion and estimate sleep from decreased movement. Participants completed a sleep diary every morning to consolidate the accelerometry data, providing information on bedtimes, wake times, and restorative sleep^96^. On day 15, the devices were returned via postal service and the data was downloaded using GENEActiv software. Currently, shared actigraphy data is restricted to measures of mean sleep onset, wake time, duration, midpoint, and restorative sleep (session 1 only). The raw actigraphy data for sessions 1 and 2 are held by Kathleen Merikangas, National Institute of Mental Health, and Ian Hickie, University of Sydney, Australia, and may be able to be shared by contacting them.

#### Early Life and Demographic Factors

Perinatal and postnatal information (e.g., maternal smoking and drinking; gestation age, birth weight, breastfeeding) were provided by the mother. Questions were adapted from earlier work^97,98^ and other freely available sources (The World Health Organisation Global Adult Tobacco Survey [GATS, 2^nd^ Edition]). Family demographic information was also provided by the parent attending session 1. These included self-report of ancestry (based on questions included in the 2016 Australian Census) and socioeconomic indexes from Census neighbourhood classifications (Socio-Economic Index for Areas [SEIFA] ^74^), occupation or education (Australian Socioeconomic Index [AUSE106] ^25^), and social status (The MacArthur Scale of Subjective Social Status^75^), as well as age at twin birth, and current height and weight.

#### Dietary Behaviour

In session 1 the attending parent completed the Australian Child and Adolescent Eating Survey (AES) ^76^ for each twin. This online food frequency questionnaire assesses the dietary intake of children and adolescents aged 2 to 17 years. It includes the frequency of consumption of 120 common foods, use of supplements, as well as eating and behaviours. Although not suitable for estimating absolute intakes, individuals can be classified into quintiles of intake for categories including total energy, protein, carbohydrate, sugars, fibre, and vitamin and mineral intake^76^. For a sub-sample of QTAB participants, these dietary intake assessments have been analysed and supplement gut microbiome measures obtained from stool samples^99^.

#### COVID-19 Interruption to Session 2 Data Collection and Pandemic Specific Assessments

Session 2 data collection, which began in November 2019, was paused on the 25^th^ of March 2020 as Brisbane entered a nation-wide shutdown (including the closure of schools). In late June 2020, session 2 data collection resumed, and continued until the conclusion of the QTAB study in January 2021. 47 families completed session 2 prior to the COVID19 interruption, and 105 families following the resumption of data collection for session 2 (see variable *lockdown_ses02* in the COVID-19 dataset (*10_covid-19*.*tsv*). There was no change to the study protocol or collection methodology post-interruption (i.e., twin and parent assessments were completed in-person at the study centre). Due to the COVID-19 interruption, the interval between session 1 and 2 was extended to a maximum of 30 months. Further, to maximise participation while constrained by funding deadlines, the minimum interval was reduced to 13 months. Preference was given to “overdue” families following the resumption of data collection; otherwise, there was no specific criteria when approaching families to return for their session 2 visit.

Pandemic specific assessments via questionnaire were acquired between August 2020 and January 2021. During this time, Brisbane schools were open and restrictions eased; however, various non-pharmaceutical interventions remained^100^. All families who participated in session 1 were contacted via email and invited to complete the questionnaire online at home (administered using Qualtrics software [Provo, UT, USA, https://www.qualtrics.com]). Those families who returned for session 2 but had not yet completed the COVID-19 questionnaire at home, completed the survey at the end of their session 2 visit. In total, we received responses from at least one family member (i.e., twin or parent) for 108 families, of which approximately half completed the questionnaire at home.

We adapted questions from multiple COVID-19 survey sources, including the Swinburne University of Technology (Melbourne) and the NIH Office of Behavioral and Social Sciences Research (OBSSR). We also assessed post-traumatic stress (Child PTSD Symptom Scale for children 8-18 years^101^), *resilience* (Brief Resilience Scale^81^; Posttraumatic Growth Inventory X^102^) and active and passive social media use (Multi-dimensional Scale of Facebook Use^78^; modified for all types of social media^77^). Resilience is implicated in mediating the development of mental illness following trauma^103^, while type of social media use (i.e., active “connection promoting” versus passive “non-connection promoting”) has been shown to mediate symptoms of anxiety and depressed mood in adolescents^77^, and active usage may be protective against negative consequences of social distancing. Further, a parent (usually the mother) reported on pandemic-related concerns and the impact on their work and home situation, personal finances, general wellbeing, and mental health – all factors that may mediate or moderate adolescent responses to the pandemic.

### Biological Samples

While we collected several biological samples, most samples have not yet been processed due to budgetary constraints; however, collection statistics (i.e., yes/no) are provided for each biological sample (see *Data Records*).

#### Salivary Sex-Steroid Collection

Participants collected a saliva sample upon minutes of wakening on the day of their visit. A saliva kit, including a collection tube, icepack, and step-by-step written instructions, was mailed in advance of the study session. The procedure was explained to the parent on the phone when booking their session visit. Participants were instructed to generate some saliva in their mouth and then slowly drool it into the tube until the desired amount was collected (i.e., 2 ml). Then, they placed it on the icepacks for transport. Upon arrival at the QTAB session, saliva samples were immediately placed on fresh ice and transferred to a -80C freezer. 99% of participants provided a saliva sample at session 1 and 100% at session 2. The long-term aim was to process the samples at the Stress Physiology Investigative Team (SPIT) laboratory at Iowa State University. Estradiol and testosterone concentrations (pg/mL) would be indexed using the Salimetrics Salivary Testosterone Enzyme Immunoassay ELISA kit.

#### Hair Cortisol

At sessions 1 and 2, a research assistant used scissors to cut 10–50 mg of hair from the posterior vertex region of the scalp (1 to 3 cm – enough to assess cortisol for the previous 3 months). Hair specimens have been stored at room temperature and have not yet been processed. Hair cortisol has recently emerged as a promising biomarker for long-term retrospective HPA activation^104^, with extremes of cortisol concentration levels (i.e., both lowest and highest) predicting depressive symptoms in adolescents^105^ and longitudinal change in concentration levels predicting later social anxiety^106^. Puberty is a period of HPA-axis plasticity, and the effects of stress on cortisol regulation may depend on developmental stage/pubertal maturation. A longer-term aim of QTAB was to track how cortisol functioning impacts the brain regions processing emotion and whether HPA reactivity interacts with pubertal stage and subsequent associations with depression. Hair cortisol data are held by associate researchers exclusively until the 30^th^ of June, 2027. After this date, data will be shared via an appropriate repository; please contact Liza van Eijk and Zoltan Sarnyai, James Cook University, after the exclusivity period ends.

#### Blood

Genomic DNA was extracted from a blood sample provided at session 1 using standard procedures. DNA samples for available twin participants were genotyped using the Infinium Global Screening Array-24 v3.0 BeadChip (Illumina, California, USA). Genotyping data is expected to be available in 2023. These data will be stored in a dedicated genotype repository under different participant identifiers. Researchers will require access approval to the QTAB non-imaging phenotypes dataset (*see Data Records - Non-Imaging Phenotype Data*) before participant identifiers can be linked between the imaging/phenotypic and genotypic datasets.

## Data Records

### Imaging Data

The QTAB imaging dataset is publicly available through the OpenNeuro data sharing platform^107^ (accession number ds004146; please use the latest version as updates may exist). The dataset is organised as per the Brain Imaging Data Structure (BIDS) specification v1.6.0^108^. BIDS specifies a hierarchical organisational format, with participant data stored under sub-folders denoting session (ses-01, ses-02) and then image modality: anat (structural), dwi (diffusion), func (functional), perf (asl), swi (susceptibility). An overview of the data record is available in Supplementary Table 1. Before sharing, all personally identifiable information was removed from the dataset, including facial features from the 3D whole-brain images (T1-weighted, T2-weighted, and FLAIR). We have made all data available, regardless of data quality (see *Technical Validation*).

Basic demographic data (age [rounded down], sex, handedness, family_id [to denote non-independent participants]) is provided at the top-level of the dataset directory in participants.tsv (variables and properties described in participants.json). Each scan is stored as a compressed NifTI file (.nii.gz) with an accompanying sidecar JSON file (.json) describing scan acquisition parameters. In addition, for the diffusion scans, gradient orientation information is provided in *\*_dwi*.*bvec and *_dwi*.*bval*, and for the asl scans, volume type information (i.e., m0scan, control, label) is provided in *\*_aslcontext*.*tsv*. For the emotional conflict and movie-watching (Partly Cloudy) tasks, event details (e.g., facial emotion responses for the emotional conflict task, condition timings for the Partly Cloudy task) are provided in *\*_task-emotionalconflict_events*.*tsv* and *_*task-partlycloudy_events*.*tsv* respectively (variables and properties described in accompanying *\*_task-emotionalconflict_events*.*json* and *\*_task-partlycloudy_events*.*json*). The defacing masks used to deface the 3D whole-brain images are provided alongside participant anatomical data in *\*_inv-2_MP2RAGE_defacemask*.*nii*.*gz*. Lastly, the *derivatives* folder at the top-level of the dataset directory contains quality checking metrics for imaging data (see *Technical Validation*) and MP2RAGE UNIT1 images with background noise removed (see *Usage Notes - MP2RAGE Acquisitions*).

### Non-Imaging Phenotype Data

The QTAB non-imaging phenotypes dataset is available through the Zenodo data sharing platform as a restricted access dataset^109^. Access requests can be submitted through the Zenodo platform. Please note, applicants are required to complete a data usage agreement (DUA)^110^ as part of the application process. To protect and assure the confidentiality and privacy of QTAB participants, all Recipients who are granted access to these data are expected to adhere to all terms of use outlined in the agreement. Access to the non-imaging phenotypic dataset will be granted to qualified investigators for appropriate use, as determined by authors Lachlan Strike and Katie McMahon.

The QTAB non-imaging phenotypes dataset contains restricted demographic data (i.e., age in months, zygosity, zygosity source, multiple birth status, birth order), as well as sex, handedness, and family_id, is provided in *participants_restricted*.*tsv* (variables and properties described in *participants_restricted*.*json*). Multiple assessments and scales of a common domain are grouped as per the phenotypic domains detailed in Table 3 (i.e., *Puberty, Cognition, Anxiety and/or Depression, Emotional and Social Behaviours, Social Support and Family Functioning, Stress, Sleep and Physical Health, Early Life and Family Demographics, Dietary Behaviour, COVID-19 Pandemic Specific Assessments*), and separated by session. Total/sum scores and individual items (where possible) are stored in a series of .tsv files, with variables and properties described in an accompanying JSON sidecar. Collection statistics (i.e., yes/no) for biological samples (blood, hair, saliva, urine, stool, baby tooth) are additionally provided. An overview of the non-imaging data record is available in Supplementary Table 2, with notations identifying the key measurement domain. Before sharing these phenotypes, we removed all participant identifiable information, including occupation, postcode, and special categories of personable data (i.e., free-response text).

## Technical Validation

### Imaging Data

3D whole-brain (T1-weighted, T2-weighted, and FLAIR) and TSE scans were visually checked and rated by one author (LTS). Scans were rated using a three-category scale (pass, warn, fail) and scan quality ratings are available in the *derivatives/visual_qc* folder. There was a higher percentage of warn and fail ratings for TSE, T2-weighted and FLAIR scans than T1-weighted scans (Fig. 2a), likely due to increased head movement associated with the acquisition of the TSE, T2-weighted, and FLAIR scans towards the end of the imaging session. The T1-weighted, T2-weighted, and FLAIR images were visually checked to ensure that facial features were successfully removed.

**Fig. 2.**
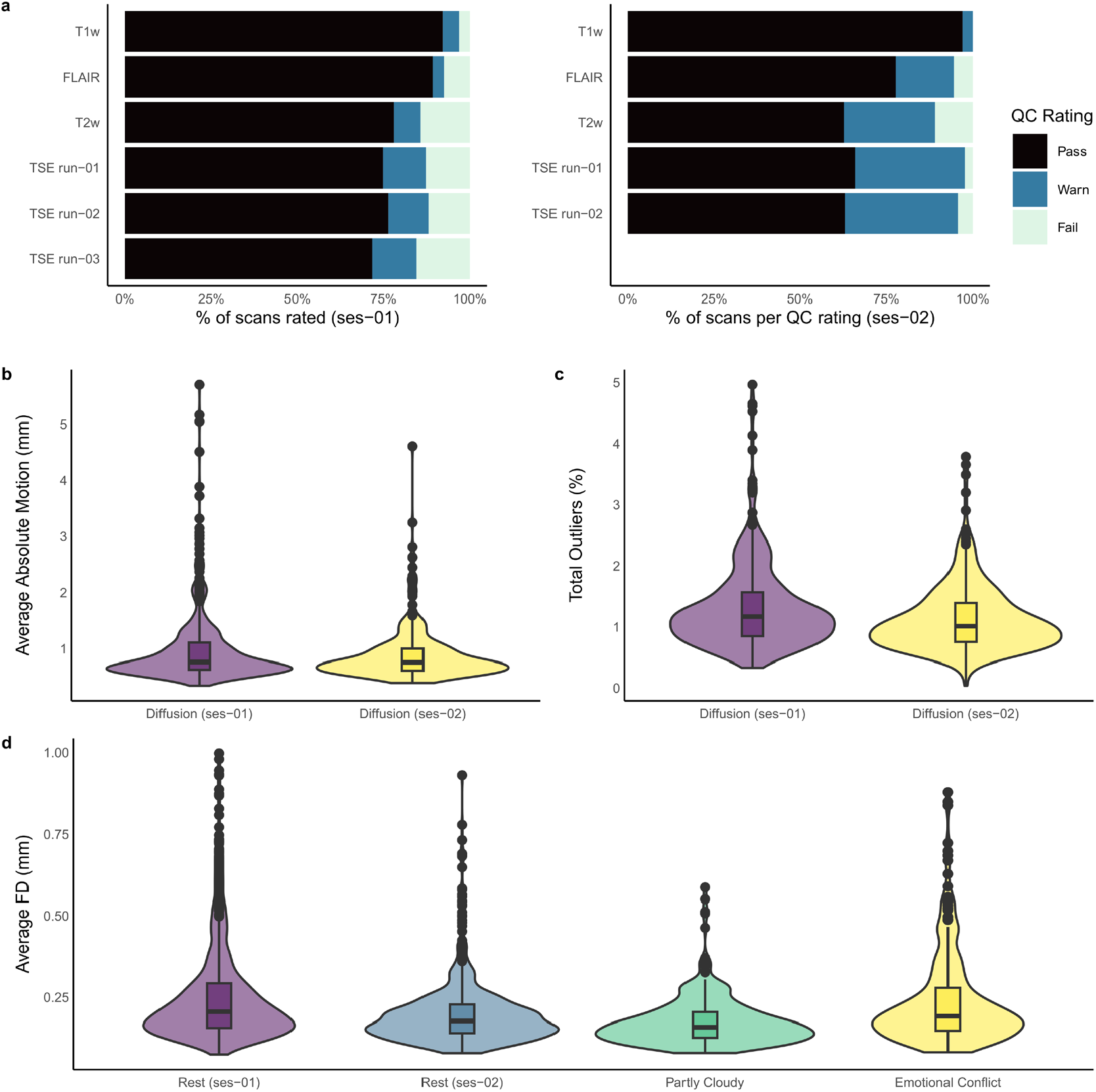
MRI image quality measures. (**a**) Visual inspection quality control (QC) ratings (pass, warn, fail) for the anatomical scans collected at session 1 and 2 (ses-01, ses-02). A *pass* rating denotes images show clear grey/white matter contrast and are free of motion or ringing artefacts. A *warn* rating indicates images show slight ringing but otherwise good grey/white matter contrast. A *fail* rating denotes images are severely impacted by motion or susceptibility artefacts with cortical and subcortical structures challenging to delineate clearly. (**b**) Violin plot showing the distribution of average absolute motion in mm (i.e., displacement relative to the first volume; lower values are better) for diffusion scans at session 1 and 2. The lower and upper hinges of the box plot represent the first quartile and third quartiles, respectively, and the horizontal line within the box represents the median. Outliers (1.5 times the interquartile range) are represented as individual data points. The median across participants of average absolute motion was similar between sessions 1 and 2 (0.77 mm and 0.76 mm, respectively). (**c**) Violin plot of the percentage of total slices classified as outliers (i.e., slices affected by severe signal dropout; lower values are better) for diffusion scans at sessions 1 and 2. The median percentage of total outlier slices was greater at session 1 (1.18%) than session 2 (1.01%). (**d**) Violin plot showing the distribution of average Framewise Displacement (FD) head motion in mm for resting-state and task fMRI scans (lower values are better); participants with mean FD greater than 1 mm not shown (rest session 1: 15 participants, rest session 2: 1 participant, emotional conflict: 4 participants). The median across participants of average FD was lowest for the Partly Cloudy movie-watching scan (0.16mm) compared to the rest and emotional conflict task scans (0.18-0.21 mm).

We used the MRtrix3 script *dwipreproc*^*111*^ (which implements the FSL tool *eddy*^112,113^) to calculate volume-to-volume motion estimates (Fig. 2b) and the percentage of detected outlier slices (i.e., slices affected by severe signal dropout; Fig. 2c); estimates available in the *derivatives/mrtrix3* folder. The median across participants of average absolute motion was 0.77 mm and 0.76 mm at sessions 1 and 2, respectively. This finding is comparable with the same metric reported in a subset of the adult HCP^114^ (median 0.83 mm). The median percentage of total outlier slices was 1.18% and 1.01% at sessions 1 and 2. The same metric was 1.89% and 0.39% in the developing/neonatal and adult HCP datasets^114^, respectively.

Image quality metrics (IQMs) were calculated for task and resting-state fMRI scans using MRIQC^115^. Framewise displacement (FD) head motion IQMs are displayed in Fig. 3d, and all IQMs are available in the *derivatives/mriqc* folder. The median across participants of average FD ranged from 0.16mm (Partly Cloudy) to 0.21mm (session 1 Rest). This finding is comparable with the same metric reported in a dataset of 3-12-year-old children^35^ (Partly Cloudy task, median = 0.29 mm) and a dataset of 8-17-year-old children^116^ (lexical processing tasks, median across all tasks and sessions = 0.17 mm). The ASL and SWI scans provided have not yet undergone quality checking or pre-processing. However, all scans were checked for apparent artefacts during the scanning session, with affected scans re-acquired (time permitting).

**Fig. 3.**
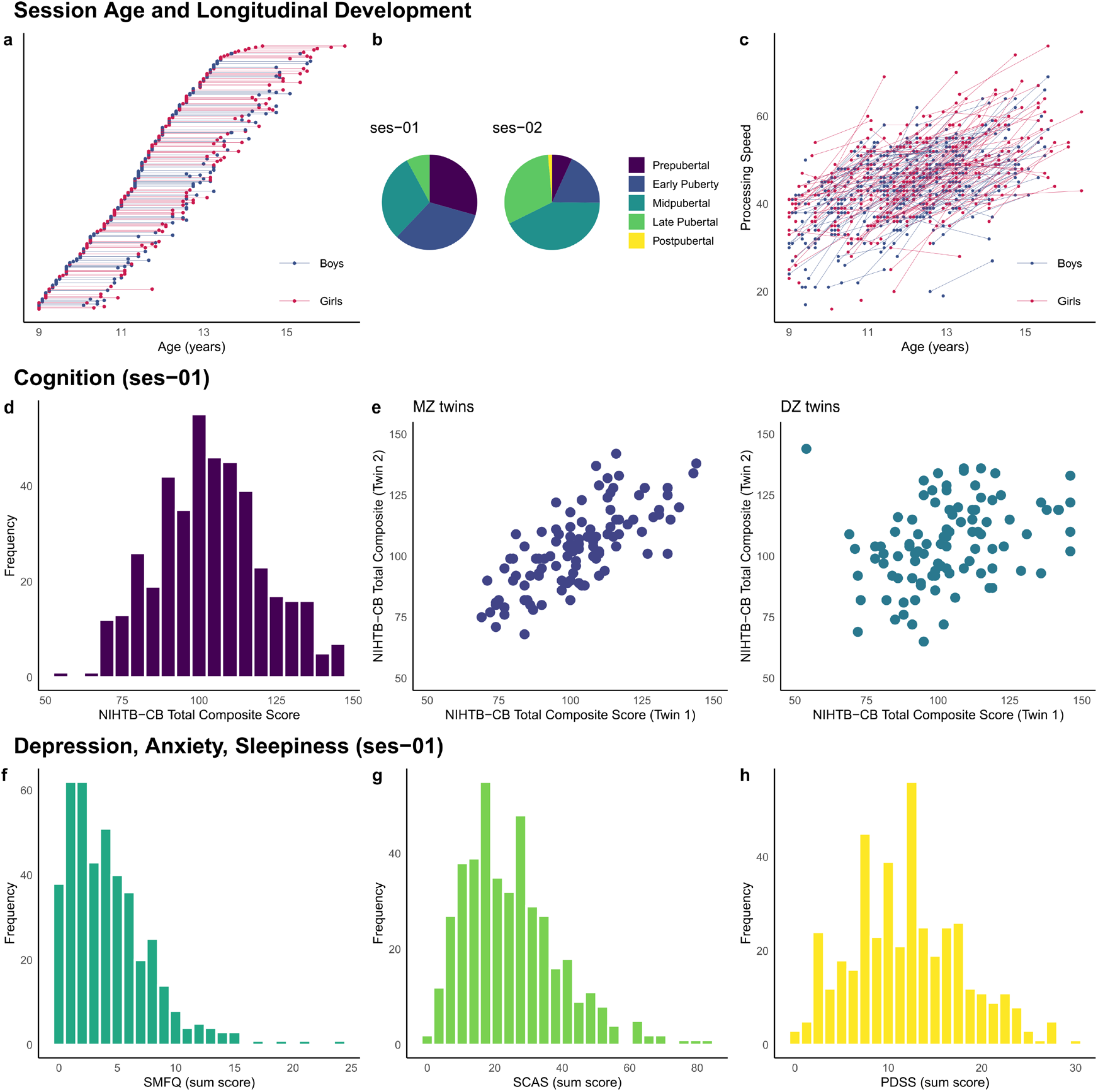
Examples supporting the technical validation of the QTAB non-imaging phenotypes. Within the 5-year funding period, the accelerated longitudinal design maximised the benefits of cross-sectional and longitudinal data collection (**a**), and developmental changes in pubertal status (**b**) and processing speed (**c**) occurred between sessions 1 and 2. The QTAB cohort was highly representative of community norms in cognitive ability (**d**), with the expected genetic relationship, i.e., identical twins were more alike than non-identical twins (**e**). In addition, the tools chosen to measure depression (**f**) and anxiety (**g**) were sensitive enough to capture individual differences and identify at-risk individuals. Variability in a depression risk factor, i.e., daytime sleepiness (**h**), was consistent with other community samples.

### Non-Imaging Phenotypes

Fig. 3 focuses on the design of QTAB and the non-imaging phenotypes. We chose an accelerated longitudinal design (Fig. 3a) so that in the first three years of the study, we worked to recruit every willing participant who was a twin between the ages of 9-14 years and who lived close to the study centre at the University of Queensland, Brisbane. We were successful in recruiting 422 twins across this age range. The long-term aim was to follow them prospectively and gather data on at least one further time point. For the 304 participants returning for session 2, the inter-session interval ranged from 1.1 to 2.5 years (M=1.7±0.3). The interval was partly influenced by pauses in data collection due to the pandemic, availability due to orthodontic treatment, and 5 years of funding. With an accelerated design^117^, as the participants age into new categories, they contribute data in every cell, i.e., the sample sizes get larger and larger as the adolescents age into these categories.

The advance of pubertal development can be seen in Fig. 3b. At session 1, more than half of the sample were classified as pre- or early-pubertal. In contrast, at session 2, approximately three-quarters of the sample had progressed to mid- or late-pubertal status, with a small number classified as post-pubertal. Developmental advances in processing speed (Fig. 3c) were also evident throughout the inter-session interval. The NIHTB-CB processing speed task records the number of correctly answered items in 85 seconds. On average, participants correctly answered an additional 8 items at session 2 compared to session 1. A trend for cross-sectional age-related increases in processing speed was also found, with older participants, in general, being able to answer more items within the given timeframe correctly.

We calculated a measure of general cognitive ability using the NIHTB-CB Total Composite Score. Age-corrected standard scores for QTAB are shown in Fig. 3d. They are very close to the US national average scores (115 and 85 indicate performance 1 SD above and below the national average of 100), with a QTAB cohort mean score of 103.5 and SD of 17.2. In addition, 7.9% of the QTAB cohort had scores of 130 or greater (top 2% based on normative NIHTB-CB data), indicating that the cohort is oversampled for high cognitive performance, and 1.4% had scores of 70 or below (bottom 2% nationally of US scores), which suggests very low cognitive functioning and may be indicative of difficulties in school or general functioning. We also found that cognitive ability is more similar between QTAB identical co-twins (MZ twins) than non-identical co-twins (DZ twins), as expected for a trait that is influenced by genetic inheritance^118^.

Distributions of symptom levels for depression and anxiety in QTAB are shown in Fig. 3f and Fig. 3g, respectively. Depression symptoms were obtained from the SMFQ, for which a cut-off of 8 has been suggested as a screen for depression in children aged 8-16 years^50^. Approximately 17% of the QTAB cohort scored 8 or higher (Fig. 3f). This is consistent with national surveys of child and adolescent health in Australia (e.g., 20% of adolescents aged 11-17 years reported experiencing high levels of psychological distress in the Young Minds Matter 2013-14 Survey^119^). SCAS sum score (Generalized anxiety) was higher in girls than boys and younger compared to older participants, consistent with other studies^49,120^. Of the 54 individuals aged 13-14 years, the mean score (M=13.3) is consistent with that reported for a community sample of 875 adolescents aged 13-14 years (M=13.5) ^49^ (sum score for males M=18.85±13.07 [versus 18.59±9.89 for QTAB]; females M=25.08±13.37 [versus 24.74±12.76 for QTAB]; combined M=21.72±13.56 [versus 21.67±11.73 for QTAB]).

Fewer symptoms were found for those aged 9-12 compared to same-age participants in Spence et al. ^120^ (M=29.7 versus 31.3 at age 9; M=26.4 versus 31.7 at age 10; M=24.0 versus 31.3 at age 11; M=22.0 versus 28.7 at age 12). Fig. 3h shows the QTAB distribution for daytime sleepiness, as assessed using the PDSS. Excessive daytime sleepiness has been associated with poor stress management and higher levels of depressive mood in adolescents aged 14-19 years^121^. A cut-off above 17 has previously been used to identify excessive sleepiness in 618 children aged 10 to 12 years^122^, identifying 18% of the sample. With the same cut-off threshold, 17% of the QTAB cohort would be classified as having excessive daytime sleepiness.

## Usage Notes

### Analysis of Twin Data

For analyses of genetic (co)variance in the QTAB dataset, we suggest investigators familiarise themselves with the classic twin model^123^. Investigators not interested in genetic (co)variance should nevertheless consider the correlated nature of twin data (i.e., the non-independence of participants) as it may violate statistical test assumptions^124^. Mixed models, which use random effects to model the correlation among twin pairs^125^, and structural equation modelling using the classic twin design^18^, are widely used approaches in controlling familial relatedness. Example code is provided online (https://github.com/QTAB-STUDY/twin-data-models).

### MP2RAGE Acquisitions

The amplified background noise in the T1w MP2RAGE uniform image (*_UNIT1) can cause registration and segmentation issues. One approach is to remove background noise from the MP2RAGE uniform image using the inversion time images^126,127^ (see *derivatives/UNIT1_denoised* for an implementation using AFNI^109^). Another technique is to input brain-extracted (i.e., skull-stripped) MP2RAGE uniform images to automated processing pipelines (e.g., FreeSurfer, fMRIPrep). We found that creating a brain mask based on the second inversion time image (*_inv-2_MP2RAGE) and applying this mask to the MP2RAGE uniform image resulted in successful brain extractions (https://github.com/QTAB-STUDY/pre-processing).

### Geometric Distortion Correction

Diffusion and rs-fMRI scans were acquired using reversed-phase encoding directions to correct geometric distortions in the images. We recommend using the FSL tools *topup* and *eddy* to correct for distortions and movement in the diffusion scans and *topup* and *applytop* to correct the rs-fMRI scans. Similarly, we recommend using the reversed-phase encoding field maps and the FSL tools *topup* and FEAT to correct the task fMRI scans. Processing pipelines implementing these tools are available online (https://github.com/QTAB-STUDY/pre-processing). Preprocessing pipelines such as fMRIPrep^128^ may also be useful for distortion correction.

### Movie-Watching Task

The video stimulus for the movie-watching task was transcoded from a PAL format DVD, resulting in a slightly faster running video than used in prior studies^34,35^. Specifically, the Partly Cloudy video used in the QTAB study was 5:31 minutes:seconds long (5:01 without end credits), compared to 5:45 minutes (5:14 without end credits) in non-PAL format. Event time codes provided in the QTAB dataset (i.e., onset and duration for mental, pain, social, and control conditions; \**_task-emotionalconflict_events*.*tsv*) correspond to the timings of a prior study^34,129^, converted to match the QTAB video stimulus. However, we recommend researchers obtain a PAL format DVD version of the film (or equivalent) to verify these condition timings (Partly Cloudy is included on physical media versions of the Disney/Pixar film Up [2009]).

### Miscellaneous Imaging

Multiple TSE scans were collected to average the TSE runs to improve image quality; see Shaw et al. ^130^ for implementation of TSE alignment and averaging using the QTAB dataset. Two participants (twin pair sub-0109 and sub-0113) required an increased number of slices for their session 1 structural scans (T1w, T2w, FLAIR) for adequate brain coverage. Three participants (twin pair sub-0200 and sub-0419, sub-0373) have a reduced number of slices in their session 1 structural scans (T1w, T2w, FLAIR) due to operator error. Four participants have task fMRI but not corresponding field map scans (sub-1207, sub-7877, sub-8742, sub-9549). One participant has an incomplete T1w acquisition at session 2 (sub-0271, missing UNIT1, inv-1 images, but has UNIT1_denoised, inv-2 images). During session 1, the T1w MP2RAGE sequence name changed (MP2RAGE_wip900C_VE11C to MP2RAGE_wip900D_VE11C); however, there was no change to the sequence parameters.

### Non-Imaging Phenotypes

Reversed scored questionnaire items have been re-coded in the non-imaging phenotypes dataset. Parent responses for several scales (i.e., APQ, FAD, LTE, PaSS, PSES, PSS, BRS, and COVID-19 Experiences and Worries) are for the family (i.e., scores are the same for co-twins within a family). Australia is a multicultural country, as reflected in the QTAB ancestry measure. This measure reflects self-perceived group identification and may differ from a person’s genetic ancestry, as obtained from genotyping. We also note that 29 participants have higher puberty scale scores at session 1 than session 2 (see the *PDS_scores_ses01_greater_ses02* variable in *01_puberty-ses01*.*tsv*). We believe this disparity reflects improved self-report puberty measurement with increased age. We suggest replacing session 01 *PDS_scale_score, PDS_category_score, Gondal_score*, and *Adrenal_score* variables with the corresponding session 02 variables. Due to copyright restrictions^131^, scale items from questionnaires are not included in the non-imaging phenotypes dataset. However, we provide detailed instructions for linking item variables to the published questionnaire items (see *00_non_imaging_phenotypes_overview*.*pdf* in the non-imaging phenotypes dataset; also provided in Supplementary Table 3). Where necessary, we contacted scale/questionnaire authors to obtain permission to use their measure and share the data collected.

## Code Availability

DICOM format MRI data was converted to a BIDS compatible dataset using HeuDiConv^132^ (v0.9.0; https://github.com/nipy/heudiconv). Facial features were removed from structural scans using BIDSonym^133^ (v0.0.5; https://github.com/peerherholz/bidsonym) and the tools *flirt* and *fslmaths* in FSL^134^ (v5.0.1; https://fsl.fmrib.ox.ac.uk/fsl/fslwiki/). Code used in data organisation and defacing is available online (https://github.com/QTAB-STUDY/dicom-to-bids). Head movement and outlier metrics for diffusion scans were calculated using the tool *dwipreproc* in MRtrix3^111^ (v3.0_RC3; https://www.mrtrix.org). Task and resting-state fMRI quality checking metrics were calculated using MRIQC^115^ (v0.16.1; https://github.com/nipreps/mriqc).

## Acknowledgements

We are forever grateful to the twins and their families for their willingness to participate in our studies. We thank Liza van Eijk, Victoria O’Callaghan, Islay Davies, Ethan Campi, Kimberley Huang, Eleanor Roga, Michael Day, Aiman Al-Najjar, Zoie Nott, Tom Shaw, Nicole Atcheson, and Sarah Daniel for data acquisition. We thank Naomi Wray for funding the collection of metabolic samples, including detailed dietary data, Ian Hickie and Kathleen Merikangas for funding and support of actigraphy data, and Sarah Medland and ENIGMA GWAS for funding genotyping. Special thanks to Anjali Henders, Leanne Wallace, Lorelle Nunn and the many laboratory assistants at the Human Studies Unit (part of the Program in Complex Trait Genomics based at the Institute of Molecular Bioscience, University of Queensland) for the processing and storage of biological samples. Thanks also to Julie Henry for helpful discussion of social cognition measures. The QTAB project was funded by the National Health and Medical Research Council (NHMRC), Australia (Project Grant ID: 1078756 to MJW), the Queensland Brain Institute, University of Queensland, and with the assistance of resources from the Centre for Advanced Imaging and the Queensland Cyber Infrastructure Foundation, University of Queensland. We acknowledge the Queensland Twin Registry (QTwin) (https://www.qimrberghofer.edu.au/study/queensland-twin-registry-study) for generously sharing database information for recruitment. Recruitment was further facilitated through access to Twins Research Australia, a national resource supported by a Centre of Research Excellence Grant (ID: 1079102) from the NHMRC. Lastly, we thank the many researchers worldwide for providing access to their assessments.

All publications of findings using data from the QTAB project should consider including the following credit and references, which should be incorporated within the “Acknowledgements” of such publication: “This research has been conducted using the QTAB project resource^107,109^”.

## Author contributions

Lachlan T. Strike – data curation, visual quality checking, pre-processing and analysis, writing – original draft.

Narelle K. Hansell - designed the data acquisition protocol, data curation, writing – original draft.

Kai-Hsiang Chuang – designed the data acquisition protocol. Jessica L. Miller – data collection, study management.

Greig I. de Zubicaray - designed the data acquisition protocol, writing - review & editing, funding acquisition.

Paul M. Thompson - designed the data acquisition protocol, writing - review & editing, funding acquisition.

Katie L. McMahon - designed the data acquisition protocol, writing - review & editing, funding acquisition.

Margaret J. Wright - designed the data acquisition protocol, writing - review & editing, funding acquisition.

