## Supplementary Materials for "The Queensland Twin Adolescent Brain Project, a longitudinal study of adolescent brain development"

| Filename | Description |
|---|---|
| **qtab** | |
| ├──.bidsignore | Files ignored by the BIDS validator |
| ├── CHANGES | Change log |
| ├── README | README file |
| ├── dataset_description.json | Brief dataset description |
| ├── participants.json | Data dictionary for participants.tsv |
| ├── participants.tsv | Participant demographics |
| ├── **derivitives** | |
| │ ├── **UNIT1_denoised** | |
| │ │ └── **sub-0001** | |
| │ │ ├── **ses-01** | |
| │ │ │ └──**anat** | |
| │ │ │ └── sub-0001_ses-01_UNIT1_unbiased_clean.nii.gz | MP2RAGE denoised uniform scan (AFNI implementation) |
| │ │ └── **ses-02** | |
| │ │ └──**anat** | |
| │ │ └── sub-0001_ses-02_UNIT1_unbiased_clean.nii.gz | MP2RAGE denoised uniform scan (AFNI implementation) |
| │ ├── **mriqc** | |
| │ │ ├── ses-01_task-rest_bold.json | Data dictionary for ses-01_task-rest_bold.tsv |
| │ │ ├── ses-01_task-rest_bold.tsv | mriqc image quality metrics for session 1 rest scans |
| │ │ ├── ses-02_task-emotionalconflict_bold.json | Data dictionary for ses-01_task-emotionalconflict_bold.json |
| │ │ ├── ses-02_task-emotionalconflict_bold.tsv | mriqc image quality metrics for session 2 emotional conflict task scan |
| │ │ ├── ses-02_task-partlycloudy_bold.json | Data dictionary for ses-02_task-partlycloudy_bold.tsv |
| │ │ ├── ses-02_task-partlycloudy_bold.tsv | mriqc image quality metrics for session 2 partly cloudy task scan |
| │ │ ├── ses-02_task-rest_bold.json | Data dictionary for ses-02_task-rest_bold.tsv |
| │ │ └── ses-02_task-rest_bold.tsv | mriqc image quality metrics for session 2 rest scans |
| │ ├── **mrtix3** | |
| │ │ ├── ses-01_dwi.json | Data dictionary for ses-01_dwi.tsv |
| │ │ ├── ses-01_dwi.tsv | EDDY QC measures for session 1 diffusion scans |
| │ │ ├── ses-02_dwi.json | Data dictionary for ses-02_dwi.tsv |
| │ │ └── ses-02_dwi.tsv | EDDY QC measures for session 2 diffusion scans |
| │ └── **visual_qc** | |
| │ ├── anat_ses-01_qc.json | Data dictionary for anat_ses-01_qc.json |
| │ ├── anat_ses-01_qc.tsv | Visual quality ratings for session 1 anatomical scans |
| │ ├── anat_ses-02_qc.json | Data dictionary for anat_ses-02_qc.json |
| │ └── anat_ses-02_qc.tsv | Visual quality ratings for session 2 anatomical scans |
| └── **sub-0001** | |
| ├── **ses-01** | |
| │ ├── **anat** | |
| │ │ ├── sub-0001_ses-01_FLAIR.json | FLAIR acquisition parameters |
| │ │ ├── sub-0001_ses-01_FLAIR.nii.gz | FLAIR scan |
| │ │ ├── sub-0001_ses-01_T2w.json | T2w acquisition parameters |
| │ │ ├── sub-0001_ses-01_T2w.nii.gz | T2w scan |
| │ │ ├── sub-0001_ses-01_T2w_TSE_run-01.json | TSE acquisition parameters |
| │ │ ├── sub-0001_ses-01_T2w_TSE_run-01.nii.gz | TSE scan |
| │ │ ├── sub-0001_ses-01_T2w_TSE_run-02.json | TSE acquisition parameters |
| │ │ ├── sub-0001_ses-01_T2w_TSE_run-02.nii.gz | TSE scan |
| │ │ ├── sub-0001_ses-01_T2w_TSE_run-03.json | TSE acquisition parameters |
| │ │ ├── sub-0001_ses-01_T2w_TSE_run-03.nii.gz | TSE scan |
| │ │ ├── sub-0001_ses-01_UNIT1.json | MP2RAGE uniform acquisition parameters |
| │ │ ├── sub-0001_ses-01_UNIT1.nii.gz | MP2RAGE uniform scan |
| │ │ ├── sub-0001_ses-01_UNIT1_denoised.json | MP2RAGE denoised uniform acquisition parameters |
| │ │ ├── sub-0001_ses-01_UNIT1_denoised.nii.gz | MP2RAGE denoised uniform scan |
| │ │ ├── sub-0001_ses-01_inv-1_MP2RAGE.json | MP2RAGE inversion time 1 acquisition parameters |
| │ │ ├── sub-0001_ses-01_inv-1_MP2RAGE.nii.gz | MP2RAGE inversion time 1 scan |
| │ │ ├── sub-0001_ses-01_inv-2_MP2RAGE.json | MP2RAGE inversion time 2 acquisition parameters |
| │ │ ├── sub-0001_ses-01_inv-2_MP2RAGE.nii.gz | MP2RAGE inversion time 2 scan |
| │ │ └── sub-0001_ses-01_inv-2_MP2RAGE_defacemask.nii.gz | Mask used to deface anatomical scans (based on inv-2_MP2RAGE) |
| │ ├── **dwi** | |
| │ │ ├── sub-0001_ses-01_dir-AP_run-01_dwi.bval | Diffusion b-value (AP direction, first acquisition) |
| │ │ ├── sub-0001_ses-01_dir-AP_run-01_dwi.bvec | Diffusion b-vector (AP direction, first acquisition) |
| │ │ ├── sub-0001_ses-01_dir-AP_run-01_dwi.json | Diffusion acquisition parameters (AP direction, first acquisition) |
| │ │ ├── sub-0001_ses-01_dir-AP_run-01_dwi.nii.gz | Diffusion scan (AP direction, first acquisition) |
| │ │ ├── sub-0001_ses-01_dir-AP_run-02_dwi.bval | Diffusion b-value (AP direction, second acquisition) |
| │ │ ├── sub-0001_ses-01_dir-AP_run-02_dwi.bvec | Diffusion b-vector (AP direction, second acquisition) |
| │ │ ├── sub-0001_ses-01_dir-AP_run-02_dwi.json | Diffusion acquisition parameters (AP direction, second acquisition) |
| │ │ ├── sub-0001_ses-01_dir-AP_run-02_dwi.nii.gz | Diffusion scan (AP direction, second acquisition) |
| │ │ ├── sub-0001_ses-01_dir-PA_run-01_dwi.bval | Diffusion b-value (PA direction, first acquisition) |
| │ │ ├── sub-0001_ses-01_dir-PA_run-01_dwi.bvec | Diffusion b-vector (PA direction, first acquisition) |
| │ │ ├── sub-0001_ses-01_dir-PA_run-01_dwi.json | Diffusion acquisition parameters (PA direction, first acquisition) |
| │ │ ├── sub-0001_ses-01_dir-PA_run-01_dwi.nii.gz | Diffusion scan (PA direction, first acquisition) |
| │ │ ├── sub-0001_ses-01_dir-PA_run-02_dwi.bval | Diffusion b-value (PA direction, second acquisition) |
| │ │ ├── sub-0001_ses-01_dir-PA_run-02_dwi.bvec | Diffusion b-vector (PA direction, second acquisition) |
| │ │ ├── sub-0001_ses-01_dir-PA_run-02_dwi.json | Diffusion acquisition parameters (PA direction, second acquisition) |
| │ │ └── sub-0001_ses-01_dir-PA_run-02_dwi.nii.gz | Diffusion scan (PA direction, second acquisition) |
| │ ├── **func** | |
| │ │ ├── sub-0001_ses-01_task-rest_dir-AP_bold.json | rs-fMRI acquisition parameters (AP direction) |
| │ │ ├── sub-0001_ses-01_task-rest_dir-AP_bold.nii.gz | rs-fMRI scan (AP direction) |
| │ │ ├── sub-0001_ses-01_task-rest_dir-PA_bold.json | rs-fMRI acquisition parameters (PA direction) |
| │ │ └── sub-0001_ses-01_task-rest_dir-PA_bold.nii.gz | rs-fMRI scan (PA direction) |
| │ ├── **perf** | |

| | |
|---|---|
| │ │ ├── sub-0001_ses-01_asl.json | ASL acquisition parameters |
| │ │ ├── sub-0001_ses-01_asl.nii.gz | ASL scan |
| │ │ ├── sub-0001_ses-01_aslcontext.tsv | Volume types of asl scan |
| │ │ ├── sub-0001_ses-01_m0scan.json | M0 acquisition parameters |
| │ │ └── sub-0001_ses-01_m0scan.nii.gz | M0 scan |
| │ └── **swi** | |
| │ ├── sub-0001_ses-01_minIP.json | Minimum intensity projection acquisition parameters |
| │ ├── sub-0001_ses-01_minIP.nii.gz | Minimum intensity projection scan |
| │ ├── sub-0001_ses-01_part-mag_GRE.json | Magnitude acquisition parameters |
| │ ├── sub-0001_ses-01_part-mag_GRE.nii.gz | Magnitude scan |
| │ ├── sub-0001_ses-01_part-phase_GRE.json | Phase acquisition parameters |
| │ ├── sub-0001_ses-01_part-phase_GRE.nii.gz | Phase scan |
| │ ├── sub-0001_ses-01_swi.json | Combined magnitude and phase acquisition parameters |
| │ └── sub-0001_ses-01_swi.nii.gz | Combined magnitude and phase scan |
| └── **ses-02** | |
| ├── **anat** | |
| │ ├── sub-0001_ses-02_FLAIR.json | FLAIR acquisition parameters |
| │ ├── sub-0001_ses-02_FLAIR.nii.gz | FLAIR scan |
| │ ├── sub-0001_ses-02_T2w.json | T2w acquisition parameters |
| │ ├── sub-0001_ses-02_T2w.nii.gz | T2w scan |
| │ ├── sub-0001_ses-02_T2w_TSE_run-01.json | TSE acquisition parameters |
| │ ├── sub-0001_ses-02_T2w_TSE_run-01.nii.gz | TSE scan |
| │ ├── sub-0001_ses-02_T2w_TSE_run-02.json | TSE acquisition parameters |
| │ ├── sub-0001_ses-02_T2w_TSE_run-02.nii.gz | TSE scan |
| │ ├── sub-0001_ses-02_UNIT1.json | MP2RAGE uniform acquisition parameters |
| │ ├── sub-0001_ses-02_UNIT1.nii.gz | MP2RAGE uniform scan |
| │ ├── sub-0001_ses-02_UNIT1_denoised.json | MP2RAGE denoised uniform acquisition parameters |
| │ ├── sub-0001_ses-02_UNIT1_denoised.nii.gz | MP2RAGE denoised uniform scan |
| │ ├── sub-0001_ses-02_inv-1_MP2RAGE.json | MP2RAGE inversion time 1 acquisition parameters |
| │ ├── sub-0001_ses-02_inv-1_MP2RAGE.nii.gz | MP2RAGE inversion time 1 scan |
| │ ├── sub-0001_ses-02_inv-2_MP2RAGE.json | MP2RAGE inversion time 2 acquisition parameters |
| │ ├── sub-0001_ses-02_inv-2_MP2RAGE.nii.gz | MP2RAGE inversion time 2 scan |
| │ └── sub-0001_ses-02_inv-2_MP2RAGE_defacemask.nii.gz | Mask used to deface anatomical scans (based on inv-2_MP2RAGE) |
| ├── **dwi** | |
| │ ├── sub-0001_ses-02_dir-AP_run-01_dwi.bval | Diffusion b-value (AP direction, first acquisition) |
| │ ├── sub-0001_ses-02_dir-AP_run-01_dwi.bvec | Diffusion b-vector (AP direction, first acquisition) |
| │ ├── sub-0001_ses-02_dir-AP_run-01_dwi.json | Diffusion acquisition parameters (AP direction, first acquisition) |
| │ ├── sub-0001_ses-02_dir-AP_run-01_dwi.nii.gz | Diffusion scan (AP direction, first acquisition) |
| │ ├── sub-0001_ses-02_dir-AP_run-02_dwi.bval | Diffusion b-value (AP direction, second acquisition) |
| │ ├── sub-0001_ses-02_dir-AP_run-02_dwi.bvec | Diffusion b-vector (AP direction, second acquisition) |
| │ ├── sub-0001_ses-02_dir-AP_run-02_dwi.json | Diffusion acquisition parameters (AP direction, second acquisition) |
| │ ├── sub-0001_ses-02_dir-AP_run-02_dwi.nii.gz | Diffusion scan (AP direction, second acquisition) |
| │ ├── sub-0001_ses-02_dir-PA_run-01_dwi.bval | Diffusion b-value (PA direction, first acquisition) |
| │ ├── sub-0001_ses-02_dir-PA_run-01_dwi.bvec | Diffusion b-vector (PA direction, first acquisition) |
| │ ├── sub-0001_ses-02_dir-PA_run-01_dwi.json | Diffusion acquisition parameters (PA direction, first acquisition) |
| │ ├── sub-0001_ses-02_dir-PA_run-01_dwi.nii.gz | Diffusion scan (PA direction, first acquisition) |
| │ ├── sub-0001_ses-02_dir-PA_run-02_dwi.bval | Diffusion b-value (PA direction, second acquisition) |
| │ ├── sub-0001_ses-02_dir-PA_run-02_dwi.bvec | Diffusion b-vector (PA direction, second acquisition) |
| │ ├── sub-0001_ses-02_dir-PA_run-02_dwi.json | Diffusion acquisition parameters (PA direction, second acquisition) |
| │ └── sub-0001_ses-02_dir-PA_run-02_dwi.nii.gz | Diffusion scan (PA direction, second acquisition) |
| ├── **fmap** | |
| │ ├── sub-0001_ses-02_dir-AP_epi.json | Field map acquisition parameters (AP direction) |
| │ ├── sub-0001_ses-02_dir-AP_epi.nii.gz | Field map scan (AP direction) |
| │ ├── sub-0001_ses-02_dir-PA_epi.json | Field map acquisition parameters (PA direction) |
| │ └── sub-0001_ses-02_dir-PA_epi.nii.gz | Field map scan (PA direction) |
| ├── **func** | |
| │ ├── sub-0001_ses-02_task-emotionalconflict_bold.json | t-fMRI acquisition parameters |
| │ ├── sub-0001_ses-02_task-emotionalconflict_bold.nii.gz | t-fMRI scan |
| │ ├── sub-0001_ses-02_task-emotionalconflict_events.json | Data dictionary for sub-0001_ses-02_task-emotionalconflict_events.json |
| │ ├── sub-0001_ses-02_task-emotionalconflict_events.tsv | t-fMRI response data |
| │ ├── sub-0001_ses-02_task-partlycloudy_bold.json | t-fMRI acquisition parameters |
| │ ├── sub-0001_ses-02_task-partlycloudy_bold.nii.gz | t-fMRI scan |
| │ ├── sub-0001_ses-02_task-partlycloudy_events.json | Data dictionary for sub-0001_ses-02_task-partlycloudy_events.tsv |
| │ ├── sub-0001_ses-02_task-partlycloudy_events.tsv | t-fMRI event data |
| │ ├── sub-0001_ses-02_task-rest_dir-AP_bold.json | rs-fMRI acquisition parameters (AP direction) |
| │ ├── sub-0001_ses-02_task-rest_dir-AP_bold.nii.gz | rs-fMRI scan (AP direction) |
| │ ├── sub-0001_ses-02_task-rest_dir-PA_bold.json | rs-fMRI acquisition parameters (PA direction) |
| │ └── sub-0001_ses-02_task-rest_dir-PA_bold.nii.gz | rs-fMRI scan (PA direction) |
| ├── **perf** | |
| │ ├── sub-0001_ses-02_asl.json | ASL acquisition parameters |
| │ ├── sub-0001_ses-02_asl.nii.gz | ASL scan |
| │ ├── sub-0001_ses-02_aslcontext.tsv | Volume types of asl scan |
| │ ├── sub-0001_ses-02_m0scan.json | M0 acquisition parameters |
| │ └── sub-0001_ses-02_m0scan.nii.gz | M0 scan |
| └── **swi** | |
| ├── sub-0001_ses-02_minIP.json | Minimum intensity projection acquisition parameters |
| ├── sub-0001_ses-02_minIP.nii.gz | Minimum intensity projection scan |
| ├── sub-0001_ses-02_part-mag_GRE.json | Magnitude acquisition parameters |
| ├── sub-0001_ses-02_part-mag_GRE.nii.gz | Magnitude scan |
| ├── sub-0001_ses-02_part-phase_GRE.json | Phase acquisition parameters |
| ├── sub-0001_ses-02_part-phase_GRE.nii.gz | Phase scan |

| | |
|---|---|
| ├── sub-0001_ses-02_swi.json | Combined magnitude and phase acquisition parameters |
| └── sub-0001_ses-02_swi.nii.gz | Combined magnitude and phase scan |

**Supplementary Table 1** Overview of the QTAB OpenNeuro dataset (imaging).
**Bold, purple font** indicates a folder.
*dir* denotes phase encoding direction (*AP* anterior-posterior or *PA* posterior-anterior).
*inv* denotes inversion time (inv-1 or inv-2).
*run* indexes multiple scans acquired in the same session using the same acquisition parameters (e.g. run-01 is the first acquisition, run-02 is the second acquisition).
*ses* denotes session (ses-01 [first session] or ses-02 [second session]).

| Filename | Description/Domain |
| --- | --- |
| **non-imaging-phenotypes** | |
| ├── 00_non_imaging_phenotypes_overview.pdf | Description of the items and scales used |
| ├── 01_puberty_ses-01.json | Data dictionary for Puberty data: session 1 |
| ├── 01_puberty_ses-01.tsv | Puberty data: session 1 |
| ├── 01_puberty_ses-02.json | Data dictionary for Puberty data: session 2 |
| ├── 01_puberty_ses-02.tsv | Puberty data: session 2 |
| ├── 02_cognition_ses-01.json | Data dictionary for Cognition data: session 1 |
| ├── 02_cognition_ses-01.tsv | Cognition data: session 1 |
| ├── 02_cognition_ses-02.json | Data dictionary for Cognition data: session 2 |
| ├── 02_cognition_ses-02.tsv | Cognition data: session 2 |
| ├── 03_anxiety_depression_ses-01.json | Data dictionary for Anxiety and/or Depression data: session 1 |
| ├── 03_anxiety_depression_ses-01.tsv | Anxiety and/or Depression data: session 1 |
| ├── 03_anxiety_depression_ses-02.json | Data dictionary for Anxiety and/or Depression data: session 2 |
| ├── 03_anxiety_depression_ses-02.tsv | Anxiety and/or Depression data: session 2 |
| ├── 04_emot_soc_behav_ses-01.json | Data dictionary for Emotional and Social Behaviours data: session 1 |
| ├── 04_emot_soc_behav_ses-01.tsv | Emotional and Social Behaviours data: session 1 |
| ├── 04_emot_soc_behav_ses-02.json | Data dictionary for Emotional and Social Behaviours data: session 2 |
| ├── 04_emot_soc_behav_ses-02.tsv | Emotional and Social Behaviours data: session 2 |
| ├── 05_social_support_family_functioning_ses-01.json | Data dictionary for Social Support and Functioning data: session 1 |
| ├── 05_social_support_family_functioning_ses-01.tsv | Social Support and Functioning data: session 1 |
| ├── 05_social_support_family_functioning_ses-02.json | Data dictionary for Social Support and Functioning data: session 2 |
| ├── 05_social_support_family_functioning_ses-02.tsv | Social Support and Functioning data: session 2 |
| ├── 06_stress_ses-01.json | Data dictionary for Stress data: session 1 |
| ├── 06_stress_ses-01.tsv | Stress data: session 1 |
| ├── 06_stress_ses-02.json | Data dictionary for Stress data: session 2 |
| ├── 06_stress_ses-02.tsv | Stress data: session 2 |
| ├── 07_sleep_physical_health_ses-01.json | Data dictionary for Sleep and Physical Health data: session 1 |
| ├── 07_sleep_physical_health_ses-01.tsv | Sleep and Physical Health data: session 1 |
| ├── 07_sleep_physical_health_ses-02.json | Data dictionary for Sleep and Physical Health data: session 2 |
| ├── 07_sleep_physical_health_ses-02.tsv | Sleep and Physical Health data: session 2 |
| ├── 08_early_life_family_demographics.json | Data dictionary for Early Life and Family Demographics data |
| ├── 08_early_life_family_demographics.tsv | Early Life and Family Demographics data |
| ├── 09_dietary_behaviour_ses-01.json | Data dictionary for Dietary Behaviour data: session 1 |
| ├── 09_dietary_behaviour_ses-01.tsv | Dietary Behaviour data: session 1 |
| ├── 10_covid19.json | Data dictionary for COVID-19 Pandemic Specific Assessments data |
| ├── 10_covid19.tsv | COVID-19 Pandemic Specific Assessments data |
| ├── 11_biological_samples.json | Data dictionary for Biological samples collection statistics |
| ├── 11_biological_samples.tsv | Biological samples collection statistics |
| ├── participants_restricted.json | Data dictionary for participants_restricted.tsv |
| └── participants_restricted.tsv | Participant demographics (age in months, zygosity, zygosity source, multiple birth status, birth order) |

**Supplementary Table 2** Overview of the QTAB Zenodo dataset (non-imaging phenotypes). Phenotypic domains (e.g., Puberty, Cognition, Anxiety and/or Depression) are detailed in Table 3 (main text).
**Bold, purple font** indicates a folder.

**Supplementary Table 3** Overview of questionnaire scale items.
Due to copyright restrictions, scale items from questionnaires are not included in the non-imaging phenotypes dataset. However, we provide detailed instructions for linking item variables to the published questionnaire items (see below). Where necessary, we contacted scale/questionnaire authors to obtain permission to use their measure and share the data collected.

### Puberty

#### Pubertal Developmental Scale (PDS)

- Carskadon, M. A., & Acebo, C. (1993). A self-administered rating scale for pubertal development. *J Adolesc Health*, *14*(3), 190-195. https://doi.org/10.1016/1054-139x(93)90004-9

NOTE: minor version variations of this scale exist.

- We have used the item response options shown in the "Download the Puberty Scale PDF" link found at **http://www.sleepforscience.org/contentmgr/showdetails.php/id/91**
- Minor wording changes were undertaken for the parent version (e.g. ***For (twin name),*** have you noticed…..)

| QTAB Variable | Scale Item Source | Comment |
|---|---|---|
| **PDS01 (self-report)**<br>**pPDS01 (parent-report)** | Carskadon et al 1993, Table 1, Question 1 | Response Option "has not yet begun to spurt" was changed to "has not yet begun to spurt or grow really fast" as some participants were unfamiliar with the word "spurt". |
| **PDS02**<br>**pPDS02** | Carskadon et al 1993, Table 1, Question 2 | |
| **PDS03**<br>**pPDS03** | Carskadon et al 1993, Table 1, Question 3 | |
| **PDS04**<br>**pPDS04** | Carskadon et al 1993, Table 1<br>Question 4 FORM FOR BOYS<br>Question 4 FORM FOR GIRLS | |
| **PDS05**<br>**pPDS05** | Carskadon et al 1993, Table 1<br>Question 5 FORM FOR BOYS<br>Question 5a FORM FOR GIRLS | |
| **PDS05b**<br>**pPDS05b** | Carskadon et al 1993, Table 1<br>Question 5b FORM FOR GIRLS | |

#### Sexual Maturation Scale (SMS)

- Morris, N. M., & Udry, J. R. (1980). Validation of a self-administered instrument to assess stage of adolescent development. *Journal of Youth and Adolescence*, *9*(3), 271-280. https://doi.org/10.1007/BF02088471

| QTAB Variable | Scale Item Source | Comment |
|---|---|---|
| **SMS01 (self-report)**<br>**pSMS01 (parent-report)** | For Girls: Morris et al 1980, Figure 1<br>For Boys: Morris et al 1980, Figure 4 | |
| **SMS02**<br>**pSMS02** | For Girls: Morris et al 1980, Figure 2<br>For Boys: Morris et al 1980, Figure 3 | |

### Cognition

#### Prospective and Retrospective Memory Questionnaire for Children (PRMQC)

- Talbot, K. D., & Kerns, K. A. (2014). Event- and time-triggered remembering: the impact of attention deficit hyperactivity disorder on prospective memory performance in children. *J Exp Child Psychol*, *127*, 126-143. https://doi.org/10.1016/j.jecp.2014.02.011

Note: gender neutral pronous used throughout

| QTAB Variable | Questionnaire Item Source | Comment |
|---|---|---|
| **pPRMQ01** | Talbot et al 2014, Appendix, 1st question | |
| **pPRMQ02** | Talbot et al 2014, Appendix, 2nd question | "he/she" replaced with "they" |
| **pPRMQ03** | Talbot et al 2014, Appendix, 3rd question | "he/she" replaced with "they";<br>"him/her" replaced with "them";<br>"his/her" replaced with "their" |
| **pPRMQ04** | Talbot et al 2014, Appendix, 4th question | "he/she" replaced with "they" |
| **pPRMQ05** | Talbot et al 2014, Appendix, 5th question | "he/she" replaced with "they" |
| **pPRMQ06** | Talbot et al 2014, Appendix, 6th question | |
| **pPRMQ07** | Talbot et al 2014, Appendix, 7th question | "his/her" replaced with "their" |
| **pPRMQ08** | Talbot et al 2014, Appendix, 8th question | "him/her" replaced with "them" |
| **pPRMQ09** | Talbot et al 2014, Appendix, 9th question | |

| | | |
|---|---|---|
| **pPRMQ10** | Talbot et al 2014, Appendix, 10th question | "him/her" replaced with "them" |
| **pPRMQ11** | Talbot et al 2014, Appendix, 11th question | "he/she" replaced with "they"; "his/her" replaced with "their" |
| **pPRMQ12** | Talbot et al 2014, Appendix, 12th question | "he/she" replaced with "they" |
| **pPRMQ13** | Talbot et al 2014, Appendix, 13th question | "he/she" replaced with "they" |
| **pPRMQ14** | Talbot et al 2014, Appendix, 14th question | "he/she" replaced with "they" |
| **pPRMQ15** | Talbot et al 2014, Appendix, 15th question | "he/she" replaced with "they" |
| **pPRMQ16** | Talbot et al 2014, Appendix, 16th question | "he/she" replaced with "they" |

### Empathy Questionnaire for Children and Adolescents (EmQue-CA)

- Overgaauw, S., Rieffe, C., Broekhof, E., Crone, E. A., & Guroglu, B. (2017). Assessing Empathy across Childhood and Adolescence: Validation of the Empathy Questionnaire for Children and Adolescents (EmQue-CA). *Front Psychol*, *8*, 870. https://doi.org/10.3389/fpsyg.2017.00870
- Variables were presented in the order shown in EmQue-CA_UK.pdf found at https://www.focusonemotions.nl/empathy-questionnaire

| QTAB Variable | Questionnaire Item Source | Comment |
|---|---|---|
| **EmQue1.1** | Overgaauw et al 2017, Table 4, Item 1.1 | |
| **EmQue1.2** | Overgaauw et al 2017, Table 4, Item 1.2 | |
| **EmQue1.3** | Overgaauw et al 2017, Table 4, Item 1.3 | |
| **EmQue1.4** | Overgaauw et al 2017, Table 4, Item 1.4 | |
| **EmQue1.5** | Overgaauw et al 2017, Table 4, Item 1.5 | |
| **EmQue1.6** | Overgaauw et al 2017, Table 4, Item 1.6 | |
| **EmQue2.1** | Overgaauw et al 2017, Table 4, Item 2.1 | |
| **EmQue2.2** | Overgaauw et al 2017, Table 4, Item 2.2 | |
| **EmQue2.3** | Overgaauw et al 2017, Table 4, Item 2.3 | "had" used instead of "has" |
| **EmQue3.1** | Overgaauw et al 2017, Table 4, Item 3.1 | "them" used instead of "him" |
| **EmQue3.2** | Overgaauw et al 2017, Table 4, Item 3.2 | |
| **EmQue3.3** | Overgaauw et al 2017, Table 4, Item 3.3 | |
| **EmQue3.4** | Overgaauw et al 2017, Table 4, Item 3.4 | |
| **EmQue3.5** | Overgaauw et al 2017, Table 4, Item 3.5 | |

**Anxiety and/or Depression**

### Spence Children's Anxiety Scale (SCAS)

- Spence, S. H., Barrett, P. M., & Turner, C. M. (2003). Psychometric properties of the Spence Children's Anxiety Scale with young adolescents. *J Anxiety Disord*, *17*(6), 605-625. https://doi.org/10.1016/s0887-6185(02)00236-0
- The parent-report version is available at https://www.scaswebsite.com/ - there are no filler items in the parent version and so numbering differs from the child self-report scale. The same items are grouped together below.

NOTE: gender neutral pronouns were used in the parent-report version at session 2 (ses-02), but not at session 1 (ses-01)

| QTAB Variable | Scale Item Source | Comment |
|---|---|---|
| **SCAS01 (Self-report)**<br>**pSCAS01 (Parent-report)** | **Self:** Spence et al 2003, Table 2, Item 1<br>**Parent:** SCAS website parent version, Item 1 | |
| **SCAS02**<br>**pSCAS02** | **Self:** Spence et al 2003, Table 2, Item 2<br>**Parent:** SCAS website parent version, Item 2 | |
| **SCAS03**<br>**pSCAS03** | **Self:** Spence et al 2003, Table 2, Item 3<br>**Parent:** SCAS website parent version, Item 3 | **Parent-report:**<br>**ses-01**"s/he" used instead of "my child"<br>**ses-02 "**s(he)" replaced with "they"; "his/her" replaced with "their" |
| **SCAS04**<br>**pSCAS04** | **Self:** Spence et al 2003, Table 2, Item 4<br>**Parent:** SCAS website parent version, Item 4 | |
| **SCAS05**<br>**pSCAS05** | **Self:** Spence et al 2003, Table 2, Item 5<br>**Parent:** SCAS website parent version, Item 5 | **Parent-report:**<br>**ses-02 "**his/her" replaced with "their" |
| **SCAS06**<br>**pSCAS06** | **Self:** Spence et al 2003, Table 2, Item 6<br>**Parent:** SCAS website parent version, Item 6 | **Parent-report:**<br>**ses-02 "**s(he)" replaced with "they" |
| **SCAS07**<br>**pSCAS07** | **Self:** Spence et al 2003, Table 2, Item 7<br>**Parent:** SCAS website parent version, Item 7 | **Parent-report:**<br>**ses-02 "**s(he)" replaced with "they" |
| **SCAS08**<br>**pSCAS08** | **Self:** Spence et al 2003, Table 2, Item 8<br>**Parent:** SCAS website parent version, Item 8 | |
| **SCAS09**<br>**pSCAS09** | **Self:** Spence et al 2003, Table 2, Item 9<br>**Parent:** SCAS website parent version, Item 9 | **Parent-report:**<br>**ses-02 "**s(he)" replaced with "they"; "him/herself" replaced with "themself" |
| **SCAS10**<br>**pSCAS10** | **Self:** Spence et al 2003, Table 2, Item 10<br>**Parent:** SCAS website parent version, Item 10 | **Parent-report:**<br>**ses-02 "**s(he)" replaced with "they" |

| | | |
|---|---|---|
| **SCAS11** | Positive filler item – self-report only<br>SCAS website child version, Item 11 | |
| **SCAS12**<br>**pSCAS11** | **Self:** Spence et al 2003, Table 2, Item 12<br>**Parent:** SCAS website parent version, Item 11 | |
| **SCAS13**<br>**pSCAS12** | **Self:** Spence et al 2003, Table 2, Item 13<br>**Parent:** SCAS website parent version, Item 12 | **Parent-report:**<br>**ses-02 "**s(he)" replaced with "they" |
| **SCAS14**<br>**pSCAS13** | **Self:** Spence et al 2003, Table 2, Item 14<br>**Parent:** SCAS website parent version, Item 13 | **Parent-report:**<br>**ses-02 "**s(he)" replaced with "they" |
| **SCAS15**<br>**pSCAS14** | **Self:** Spence et al 2003, Table 2, Item 15<br>**Parent:** SCAS website parent version, Item 14 | **Parent-report:**<br>**ses-02 "**s(he)" replaced with "they"; "his/her" replaced with "their" |
| **SCAS16**<br>**pSCAS15** | **Self:** Spence et al 2003, Table 2, Item 16<br>**Parent:** SCAS website parent version, Item 15 | **Parent-report:**<br>**ses-02 "**s(he)" replaced with "they" |
| **SCAS17** | Positive filler item – self-report only<br>SCAS website child version, Item 17 | |
| **SCAS18**<br>**pSCAS16** | **Self:** Spence et al 2003, Table 2, Item 18<br>**Parent:** SCAS website parent version, Item 16 | |
| **SCAS19**<br>**pSCAS17** | **Self:** Spence et al 2003, Table 2, Item 19<br>**Parent:** SCAS website parent version, Item 17 | **Parent-report:**<br>**ses-02 "**his/her" replaced with "their" |
| **SCAS20**<br>**pSCAS18** | **Self:** Spence et al 2003, Table 2, Item 20<br>**Parent:** SCAS website parent version, Item 18 | **Parent-report:**<br>**ses-02 "**s(he)" replaced with "they"; "his/her" replaced with "their" |
| **SCAS21**<br>**pSCAS19** | **Self:** Spence et al 2003, Table 2, Item 21<br>**Parent:** SCAS website parent version, Item 19 | |
| **SCAS22**<br>**pSCAS20** | **Self:** Spence et al 2003, Table 2, Item 22<br>**Parent:** SCAS website parent version, Item 20 | **Parent-report:**<br>**ses-02 "**him/her" replaced with "them" |
| **SCAS23**<br>**pSCAS21** | **Self:** Spence et al 2003, Table 2, Item 23<br>**Parent:** SCAS website parent version, Item 21 | |
| **SCAS24**<br>**pSCAS22** | **Self:** Spence et al 2003, Table 2, Item 24<br>**Parent:** SCAS website parent version, Item 22 | **Parent-report:**<br>**ses-02 "**s(he)" replaced with "they" |
| **SCAS25**<br>**pSCAS23** | **Self:** Spence et al 2003, Table 2, Item 25<br>**Parent:** SCAS website parent version, Item 23 | |
| **SCAS26** | Positive filler item – self-report only<br>SCAS website child version, Item 26 | |
| **SCAS27**<br>**pSCAS24** | **Self:** Spence et al 2003, Table 2, Item 27<br>**Parent:** SCAS website parent version, Item 24 | |
| **SCAS28**<br>**pSCAS25** | **Self:** Spence et al 2003, Table 2, Item 28<br>**Parent:** SCAS website parent version, Item 25 | **Parent-report:**<br>**ses-02 "**s(he)" replaced with "they" |
| **SCAS29**<br>**pSCAS26** | **Self:** Spence et al 2003, Table 2, Item 29<br>**Parent:** SCAS website parent version, Item 26 | **Parent-report:**<br>**ses-02 "**him/her" replaced with "them" |
| **SCAS30**<br>**pSCAS27** | **Self:** Spence et al 2003, Table 2, Item 30<br>**Parent:** SCAS website parent version, Item 27 | |
| **SCAS31** | Positive filler item -self-report only<br>SCAS website child version, Item 31 | |
| **SCAS32**<br>**pSCAS28** | **Self:** Spence et al 2003, Table 2, Item 32<br>**Parent:** SCAS website parent version, Item 28 | |
| **SCAS33**<br>**pSCAS29** | **Self:** Spence et al 2003, Table 2, Item 33<br>**Parent:** SCAS website parent version, Item 29 | |
| **SCAS34**<br>**pSCAS30** | **Self:** Spence et al 2003, Table 2, Item 34<br>**Parent:** SCAS website parent version, Item 30 | |
| **SCAS35**<br>**pSCAS31** | **Self:** Spence et al 2003, Table 2, Item 35<br>**Parent:** SCAS website parent version, Item 31 | **Parent-report:**<br>**ses-02 "**s(he)" replaced with "they" |
| **SCAS36**<br>**pSCAS32** | **Self:** Spence et al 2003, Table 2, Item 36<br>**Parent:** SCAS website parent version, Item 32 | **Parent-report:**<br>**ses-02 "**his/her" replaced with "their" |
| **SCAS37**<br>**pSCAS33** | **Self:** Spence et al 2003, Table 2, Item 37<br>**Parent:** SCAS website parent version, Item 33 | **Parent-report:**<br>**ses-02 "**s(he)" replaced with "they" |
| **SCAS38** | Positive filler item – self-report only<br>SCAS website child version, Item 38 | |
| **SCAS39**<br>**pSCAS34** | **Self:** Spence et al 2003, Table 2, Item 39<br>**Parent:** SCAS website parent version, Item 34 | |
| **SCAS40**<br>**pSCAS35** | **Self:** Spence et al 2003, Table 2, Item 40<br>**Parent:** SCAS website parent version, Item 35 | **Parent-report:**<br>**ses-02 "**his/her" replaced with "their" |
| **SCAS41**<br>**pSCAS36** | **Self:** Spence et al 2003, Table 2, Item 41<br>**Parent:** SCAS website parent version, Item 36 | **Parent-report:**<br>**ses-02 "**his/her" replaced with "their" |
| **SCAS42**<br>**pSCAS37** | **Self:** Spence et al 2003, Table 2, Item 42<br>**Parent:** SCAS website parent version, Item 37 | |

| | | |
|---|---|---|
| **SCAS43** | Positive filler item – self-report only<br>SCAS website child version, Item 43 | |
| **SCAS44**<br>**pSCAS38** | **Self:** Spence et al 2003, Table 2, Item 44<br>**Parent:** SCAS website parent version, Item 38 | **Parent-report:**<br>**ses-02** "s(he)" replaced with "they" |

### Short Moods and Feelings Questionnaire (SMFQ)

- Angold, A., Costello, E. J., & Messer, S. C. (1995). Development of a short questionnaire for use in epidemiological studies of depression in children and adolescents. *Int J Methods Psychiatr Res*, *5*, 237-249.
- For free download versions of the questionnaire see: **https://devepi.duhs.duke.edu/measures/the-mood-and-feelings-questionnaire-mfq/**

NOTE: gender neutral pronouns were used in the parent-report version at session 2 (ses-02), but not at session 1 (ses-01)

| QTAB Variable | Questionnaire Item Source | Comment |
|---|---|---|
| **SMFQ01 (Self-report)**<br>**pSMFQ01 (Parent-report)** | Website Child **Self-Report** Short Version, Item 1<br>Website **Parent-Report** on Child Short Version, Item 1 | **Parent-report:**<br>**ses-02** "s/he" replaced with "they" |
| **SMFQ02**<br>**pSMFQ02** | Website Child **Self-Report** Short Version, Item 2<br>Website **Parent-Report** on Child Short Version, Item 2 | **Parent-report:**<br>**ses-02** "s/he" replaced with "they" |
| **SMFQ03**<br>**pSMFQ03** | Website Child **Self-Report** Short Version, Item 3<br>Website **Parent-Report** on Child Short Version, Item 3 | **Parent-report:**<br>**ses-02** "s/he" replaced with "they" |
| **SMFQ04**<br>**pSMFQ04** | Website Child **Self-Report** Short Version, Item 4<br>Website **Parent-Report** on Child Short Version, Item 4 | **Parent-report:**<br>**ses-02** "s/he" replaced with "they" |
| **SMFQ05**<br>**pSMFQ05** | Website Child **Self-Report** Short Version, Item 5<br>Website **Parent-Report** on Child Short Version, Item 5 | **Parent-report:**<br>**ses-02** "s/he" replaced with "they" |
| **SMFQ06**<br>**pSMFQ06** | Website Child **Self-Report** Short Version, Item 6<br>Website **Parent-Report** on Child Short Version, Item 6 | **Parent-report:**<br>**ses-02** "s/he" replaced with "they" |
| **SMFQ07**<br>**pSMFQ07** | Website Child **Self-Report** Short Version, Item 7<br>Website **Parent-Report** on Child Short Version, Item 7 | **Parent-report:**<br>**ses-02** "s/he" replaced with "they" |
| **SMFQ08**<br>**pSMFQ08** | Website Child **Self-Report** Short Version, Item 8<br>Website **Parent-Report** on Child Short Version, Item 8 | **Parent-report:**<br>**ses-02** "s/he" replaced with "they" |
| **SMFQ09**<br>**pSMFQ09** | Website Child **Self-Report** Short Version, Item 9<br>Website **Parent-Report** on Child Short Version, Item 9 | **Parent-report:**<br>**ses-02** "s/he" replaced with "they"; "himself/herself" replaced with "themself" |
| **SMFQ10**<br>**pSMFQ10** | Website Child **Self-Report** Short Version, Item 10<br>Website **Parent-Report** on Child Short Version, Item 10 | **Parent-report:**<br>**ses-02** "s/he" replaced with "they" |
| **SMFQ11**<br>**pSMFQ11** | Website Child **Self-Report** Short Version, Item 11<br>Website **Parent-Report** on Child Short Version, Item 11 | **Parent-report:**<br>**ses-02** "s/he" replaced with "they"; "him/her" replaced with "them" |
| **SMFQ12**<br>**pSMFQ12** | Website Child **Self-Report** Short Version, Item 12<br>Website **Parent-Report** on Child Short Version, Item 12 | **Parent-report:**<br>**ses-02** "s/he" replaced with "they" |
| **SMFQ13**<br>**pSMFQ13** | Website Child **Self-Report** Short Version, Item 13<br>Website **Parent-Report** on Child Short Version, Item 13 | **Parent-report:**<br>**ses-02** "s/he" replaced with "they" |

### Somatic and Psychological Health Report (SPHERE-21)

- Couvy-Duchesne, B., Davenport, T. A., Martin, N. G., Wright, M. J., & Hickie, I. B. (2017). Validation and psychometric properties of the Somatic and Psychological HEalth REport (SPHERE) in a young Australian-based population sample using non-parametric item response theory. *BMC Psychiatry*, *17*(1), 279. https://doi.org/10.1186/s12888-017-1420-1
- Hickie, I. B., Davenport, T. A., Hadzi-Pavlovic, D., Koschera, A., Naismith, S. L., Scott, E. M., & Wilhelm, K. A. (2001). Development of a simple screening tool for common mental disorders in general practice. *Med J Aust*, *175*(S1), S10-17. https://doi.org/10.5694/j.1326-5377.2001.tb143784.x

| QTAB Variable | Report Item Source | Comment |
|---|---|---|
| **SPHERE02** | Couvy-Duchesne et al 2017, Fig 9, Item 2 | |
| **SPHERE03** | Couvy-Duchesne et al 2017, Fig 9, Item 3 | |
| **SPHERE05** | Couvy-Duchesne et al 2017, Fig 9, Item 5 | |
| **SPHERE07** | Couvy-Duchesne et al 2017, Fig 9, Item 7 | |
| **SPHERE08** | Couvy-Duchesne et al 2017, Fig 9, Item 8 | |
| **SPHERE12** | Couvy-Duchesne et al 2017, Fig 9, Item 12 | |
| **SPHERE15** | Couvy-Duchesne et al 2017, Fig 9, Item 15 | |

| | | |
|---|---|---|
| **SPHERE17** | Couvy-Duchesne et al 2017, Fig 9, Item 17 | |
| **SPHERE20** | Couvy-Duchesne et al 2017, Fig 9, Item 20 | |
| **SPHERE22** | Couvy-Duchesne et al 2017, Fig 9, Item 22 | |
| **SPHERE23** | Couvy-Duchesne et al 2017, Fig 9, Item 23 | |
| **SPHERE25** | Couvy-Duchesne et al 2017, Fig 9, Item 25 | |
| **SPHERE26** | Couvy-Duchesne et al 2017, Fig 9, Item 26 | |
| **SPHERE27** | Couvy-Duchesne et al 2017, Fig 9, Item 27 | |
| **SPHERE28** | Couvy-Duchesne et al 2017, Fig 9, Item 28 | |
| **SPHERE29** | Couvy-Duchesne et al 2017, Fig 9, Item 29 | We used a longer version of this item "Feeling tired after rest or relaxation?" |
| **SPHERE30** | Couvy-Duchesne et al 2017, Fig 9, Item 30 | |
| **SPHERE31** | Couvy-Duchesne et al 2017, Fig 9, Item 31 | |
| **SPHERE32** | Couvy-Duchesne et al 2017, Fig 9, Item 32 | |
| **SPHERE33** | Couvy-Duchesne et al 2017, Fig 9, Item 33 | |
| **SPHERE34** | Couvy-Duchesne et al 2017, Fig 9, Item 34 | |

**Emotional and Social Behaviours**

### Short UPPS-P Impulsive Behaviours Scale in Children (IBS)

• Geurten M, et al. Measuring Impulsivity in Children: Adaptation and Validation of a Short Version of the UPPS-P Impulsive Behaviors Scale in Children and Investigation of its Links With ADHD. J Atten Disord. 2021;25(1):105-14.

| QTAB Variable | Scale Item Source | Comment |
|---|---|---|
| **IBS01** | Geurten et al 2021, Appendix A, Item 1 | |
| **IBS02** | Geurten et al 2021, Appendix A, Item 2 | |
| **IBS03** | Geurten et al 2021, Appendix A, Item 3 | |
| **IBS04** | Geurten et al 2021, Appendix A, Item 4 | |
| **IBS05** | Geurten et al 2021, Appendix A, Item 5 | |
| **IBS06** | Geurten et al 2021, Appendix A, Item 6 | |
| **IBS07** | Geurten et al 2021, Appendix A, Item 7 | |
| **IBS08** | Geurten et al 2021, Appendix A, Item 8 | |
| **IBS09** | Geurten et al 2021, Appendix A, Item 9 | |
| **IBS10** | Geurten et al 2021, Appendix A, Item 10 | |
| **IBS11** | Geurten et al 2021, Appendix A, Item 11 | |
| **IBS12** | Geurten et al 2021, Appendix A, Item 12 | |
| **IBS13** | Geurten et al 2021, Appendix A, Item 13 | |
| **IBS14** | Geurten et al 2021, Appendix A, Item 14 | |
| **IBS15** | Geurten et al 2021, Appendix A, Item 15 | "really" was inadvertently left out |
| **IBS16** | Geurten et al 2021, Appendix A, Item 16 | |
| **IBS17** | Geurten et al 2021, Appendix A, Item 17 | |
| **IBS18** | Geurten et al 2021, Appendix A, Item 18 | |
| **IBS19** | Geurten et al 2021, Appendix A, Item 19 | |
| **IBS20** | Geurten et al 2021, Appendix A, Item 20 | |

### Strength and Difficulties Questionnaire (SDQ)

- Goodman, R. (1997). The Strengths and Difficulties Questionnaire: a research note. *J Child Psychol Psychiatry*, *38*(5), 581-586. https://doi.org/10.1111/j.1469-7610.1997.tb01545.x
- Alternate versions of the SDQ can be found at **https://www.sdqinfo.org/py/sdqinfo/b0.py** – including an "English (Austral)" version for parents or teachers of 11-17 year olds (SDQ_English(Austral)_pt11-17single.pdf)

| QTAB Variable | Questionnaire Item Source | Comment |
|---|---|---|
| **pSDQ01** | Goodman et al 1997, Appendix A, 1st item | |
| **pSDQ02** | Goodman et al 1997, Appendix A, 2nd item | |
| **pSDQ03** | Goodman et al 1997, Appendix A, 3rd item | Minor version difference: "for example, toys, treats, pencils" used instead of "(treats, toys, pencils etc)" |
| **pSDQ04** | Goodman et al 1997, Appendix A, 4th item | www.sdqinfo.org Item version used: SDQ_English(Austral)_pt11-17single |
| **pSDQ05** | Goodman et al 1997, Appendix A, 5th item | |
| **pSDQ06** | Goodman et al 1997, Appendix A, 6th item | |
| **pSDQ07** | Goodman et al 1997, Appendix A, 7th item | |
| **pSDQ08** | Goodman et al 1997, Appendix A, 8th item | |
| **pSDQ09** | Goodman et al 1997, Appendix A, 9th item | |
| **pSDQ10** | Goodman et al 1997, Appendix A, 10th item | |
| **pSDQ11** | Goodman et al 1997, Appendix A, 11th item | |
| **pSDQ12** | Goodman et al 1997, Appendix A, 12th item | www.sdqinfo.org Item version used: SDQ_English(Austral)_pt11-17single |
| **pSDQ13** | Goodman et al 1997, Appendix A, 13th item | www.sdqinfo.org Item version used: SDQ_English(Austral)_pt11-17single |

| | | |
|---|---|---|
| **pSDQ14** | Goodman et al 1997, Appendix A, 14th item | |
| **pSDQ15** | Goodman et al 1997, Appendix A, 15th item | |
| **pSDQ16** | Goodman et al 1997, Appendix A, 16th item | |
| **pSDQ17** | Goodman et al 1997, Appendix A, 17th item | |
| **pSDQ18** | Goodman et al 1997, Appendix A, 18th item | www.sdqinfo.org Item version used: SDQ_English(Austral)_pt11-17single |
| **pSDQ19** | Goodman et al 1997, Appendix A, 19th item | |
| **pSDQ20** | Goodman et al 1997, Appendix A, 20th item | |
| **pSDQ21** | Goodman et al 1997, Appendix A, 21th item | |
| **pSDQ22** | Goodman et al 1997, Appendix A, 22th item | |
| **pSDQ23** | Goodman et al 1997, Appendix A, 23h item | |
| **pSDQ24** | Goodman et al 1997, Appendix A, 24th item | |
| **pSDQ25** | Goodman et al 1997, Appendix A, 25th item | www.sdqinfo.org Item version used: SDQ_English(Austral)_pt11-17single |

### Australian-adapted Hierarchical Personality Inventory for Children (HiPIC-A)

**Also known as the Australian Child Personality Inventory (ACPI)**

- Watt, D., Hopkinson, L., Costello, S., & Roodenburg, J. (2017). Initial Validation and Refinement of the Hierarchical Inventory of Personality for Children in the Australian Context. *Australian Psychologist*, *52*(1), 61-71. https://doi.org/10.1111/ap.12213
- Hopkinson, L., Watt, D., & Roodenburg, J. (2014). Australian Validation of the Hierarchical Personality Inventory for Children (HiPIC). *The Australian Educational and Developmental Psychologist*, *31*(02), 113-124. https://doi.org/10.1017/edp.2014.3

**Adapted from:**

- Mervielde, I., & De Fruyt, F. (1999). Construction of the Hierarchical Personality Inventory for Children (HiPIC). In I. Mervielde, I. Deary, F. De Fruyt, & F. Ostendorf (Eds.), *Personality psychology in Europe. Proceedings of the Eighth European Conference on Personality Psychology* (pp. 107-111). Tilberg University Press.

NOTE: Scale items are not publicly available (for further information contact ****). ***Any use of the data must include the above references.***

- Gender neutral pronouns were used for all items.
- The QTAB data available are mean domains (factors) and facet scores (range 1-5) as shown below.

| QTAB Variable | Domain/Facet Reference | Comment |
|---|---|---|
| **pHiPICA_amen_score** | Domain: Amenability<br>Watt et al 2017, Table 2, 1st Domain | |
| **pHiPICA_cons_score** | Domain: Conscientiousness<br>Watt et al 2017, Table 2, 2nd Domain | |
| **pHiPICA_emos_score** | Domain: Emotional Stability<br>Watt et al 2017, Table 2, 3rd Domain | |
| **pHiPICA_imag_score** | Domain: Imagination<br>Watt et al 2017, Table 2, 4th Domain | |
| **pHiPICA_extr_score** | Domain: Extraversion<br>Watt et al 2017, Table 2, 5th Domain | |
| **pHiPICA_amenA1_score** | Amenability Facet: Irritability<br>Watt et al 2017, Table 2, A1 | |
| **pHiPICA_amenA2_score** | Amenability Facet: Egocentrism<br>Watt et al 2017, Table 2, A2 | |
| **pHiPICA_amenA3_score** | Amenability Facet: Compliance<br>Watt et al 2017, Table 2, A3 | |
| **pHiPICA_amenA4_score** | Amenability Facet: Dominance<br>Watt et al 2017, Table 2, A4 | |
| **pHiPICA_consC1_score** | Conscientiousness Facet: Order<br>Watt et al 2017, Table 2, C1 | |
| **pHiPICA_consC2_score** | Conscientiousness Facet: Achievement/Motivation<br>Watt et al 2017, Table 2, C2 | |
| **pHiPICA_consC3_score** | Conscientiousness Facet: Perseverance<br>Watt et al 2017, Table 2, C3 | |
| **pHiPICA_consC4_score** | Conscientiousness Facet: Concentration<br>Watt et al 2017, Table 2, C4 | |
| **pHiPICA_emosS1_score** | Emotional Stability Facet: Anxiety<br>Watt et al 2017, Table 2, S1 | |
| **pHiPICA_emosS2_score** | Emotional Stability Facet: Self confidence<br>Watt et al 2017, Table 2, S2 | |
| **pHiPICA_imagI1_score** | Imagination Facet: Creativity<br>Watt et al 2017, Table 2, I1 | |
| **pHiPICA_imagI2_score** | Imagination Facet: Curiosity<br>Watt et al 2017, Table 2, I2 | |
| **pHiPICA_imagI3_score** | Imagination Facet: Intellect<br>Watt et al 2017, Table 2, I3 | |
| **pHiPICA_extrE1_score** | Extraversion Facet: Energy | |

| | |
|---|---|
| | Watt et al 2017, Table 2, E1 |
| **pHiPICA_extrE2_score** | Extraversion Facet: Expressivity<br>Watt et al 2017, Table 2, E2 |
| **pHiPICA_extrE3_score** | Extraversion Facet: Shyness<br>Watt et al 2017, Table 2, E3 |
| **pHiPICA_extrE4_score** | Extraversion Facet: Optimism<br>Watt et al 2017, Table 2, E4 |

### Autism Spectrum Quotient – 10 items (AQ-10) (Adolescent)

- Allison, C., Auyeung, B., & Baron-Cohen, S. (2012). Toward brief "Red Flags" for autism screening: The Short Autism Spectrum Quotient and the Short Quantitative Checklist for Autism in toddlers in 1,000 cases and 3,000 controls [corrected]. *J Am Acad Child Adolesc Psychiatry*, *51*(2), 202-212 e207. https://doi.org/10.1016/j.jaac.2011.11.003

NOTE: gender neutral pronouns were used.

| QTAB Variable | Scale Item Source | Comment |
|---|---|---|
| **pAQ01** | Allison et al 2012, Table 2, AQ Adolescent, 1st Item | "S/he notices" replaced with "they notice" |
| **pAQ02** | Allison et al 2012, Table 2, AQ Adolescent, 2nd Item | "S/he usually concentrates" replaced with "they usually concentrate" |
| **pAQ03** | Allison et al 2012, Table 2, AQ Adolescent, 3rd Item | "s/he" replaced with "they" |
| **pAQ04** | Allison et al 2012, Table 2, AQ Adolescent, 4th Item | "s/he" replaced with "they" |
| **pAQ05** | Allison et al 2012, Table 2, AQ Adolescent, 5th Item | "S/he frequently finds" replaced with "they frequently find"; "s/he doesn't" replaced with "they don't" |
| **pAQ06** | Allison et al 2012, Table 2, AQ Adolescent, 6th Item | "s/he is" replaced with "they are" |
| **pAQ07** | Allison et al 2012, Table 2, AQ Adolescent, 7th Item | "s/he was" replaced with "they were"; "s/he" replaced "they. |
| **pAQ08** | Allison et al 2012, Table 2, AQ Adolescent, 8th Item | "s/he finds" replaced with "they find" |
| **pAQ09** | Allison et al 2012, Table 2, AQ Adolescent, 9th Item | "s/he finds" replaced with "they find" |
| **pAQ10** | Allison et al 2012, Table 2, AQ Adolescent, 10th Item | "s/he finds" replaced with "they find" |

### Children's Response Styles Questionnaire – rumination subscale (CRSQ)

- Abela, J. R., Brozina, K., & Haigh, E. P. (2002). An examination of the response styles theory of depression in third- and seventh-grade children: a short-term longitudinal study. *J Abnorm Child Psychol*, *30*(5), 515-527. https://www.ncbi.nlm.nih.gov/pubmed/12403154
- Abela, J. R. Z., Aydin, C. M., & Auerbach, R. P. (2007). Responses to depression in children: reconceptualizing the relation among response styles. *J Abnorm Child Psychol*, *35*, 913-927. https://doi.org/10.1007/s10802-007-9143-2

NOTE: The CRSQ consists of 25 items. Data were only collected for the 13 items of the Ruminative Response subscale, which was scored as described in Abela et al 2002.

| QTAB Variable | Questionnaire Item Source | Comment |
|---|---|---|
| **CRSQ01** | Abela et al 2007, Table 1, Item 1 | |
| **CRSQ03** | Abela et al 2007, Table 1, Item 3 | |
| **CRSQ05** | Abela et al 2007, Table 1, Item 5 | |
| **CRSQ07** | Abela et al 2007, Table 1, Item 7 | |
| **CRSQ09** | Abela et al 2007, Table 1, Item 9 | |
| **CRSQ11** | Abela et al 2007, Table 1, Item 11 | |
| **CRSQ13** | Abela et al 2007, Table 1, Item 13 | |
| **CRSQ15** | Abela et al 2007, Table 1, Item 15 | |
| **CRSQ17** | Abela et al 2007, Table 1, Item 17 | |
| **CRSQ19** | Abela et al 2007, Table 1, Item 19 | |
| **CRSQ21** | Abela et al 2007, Table 1, Item 21 | |
| **CRSQ23** | Abela et al 2007, Table 1, Item 23 | |
| **CRSQ25** | Abela et al 2007, Table 1, Item 25 | |

### Children's Attributional Style Questionnaire – Revised (CASQ-R)

- Thompson, M., Kaslow, N. J., Weiss, B., & Nolen-Hoeksema, S. (1998). Children's Attributional Style Questionnaire Revised: Psychometric examination. *Psychological Assessment*, *10*(2), 166-170. https://doi.org/10.1037/1040-3590.10.2.166

| QTAB Variable | Questionnaire Item Source | Comment |
|---|---|---|

| | | |
|---|---|---|
| **CASQ01** | Thompson et al 1998, Appendix, Item 1 | |
| **CASQ02** | Thompson et al 1998, Appendix, Item 2 | |
| **CASQ03** | Thompson et al 1998, Appendix, Item 3 | |
| **CASQ04** | Thompson et al 1998, Appendix, Item 4 | |
| **CASQ05** | Thompson et al 1998, Appendix, Item 5 | |
| **CASQ06** | Thompson et al 1998, Appendix, Item 6 | |
| **CASQ07** | Thompson et al 1998, Appendix, Item 7 | |
| **CASQ08** | Thompson et al 1998, Appendix, Item 8 | |
| **CASQ09** | Thompson et al 1998, Appendix, Item 9 | |
| **CASQ10** | Thompson et al 1998, Appendix, Item 10 | |
| **CASQ11** | Thompson et al 1998, Appendix, Item 11 | |
| **CASQ12** | Thompson et al 1998, Appendix, Item 12 | |
| **CASQ13** | Thompson et al 1998, Appendix, Item 13 | |
| **CASQ14** | Thompson et al 1998, Appendix, Item 14 | |
| **CASQ15** | Thompson et al 1998, Appendix, Item 15 | |
| **CASQ16** | Thompson et al 1998, Appendix, Item 16 | |
| **CASQ17** | Thompson et al 1998, Appendix, Item 17 | |
| **CASQ18** | Thompson et al 1998, Appendix, Item 18 | |
| **CASQ19** | Thompson et al 1998, Appendix, Item 19 | |
| **CASQ20** | Thompson et al 1998, Appendix, Item 20 | “doing” was inadvertently left out (sentence meaning essentially unchanged) |
| **CASQ21** | Thompson et al 1998, Appendix, Item 21 | |
| **CASQ22** | Thompson et al 1998, Appendix, Item 22 | |
| **CASQ23** | Thompson et al 1998, Appendix, Item 23 | |
| **CASQ24** | Thompson et al 1998, Appendix, Item 24 | |

### Early Adolescent Temperament Questionnaire – Revised (EATQ-R)

- Oldehinkel, A. J., Hartman, C. A., De Winter, A. F., Veenstra, R., & Ormel, J. (2004). Temperament profiles associated with internalizing and externalizing problems in preadolescence. *Dev Psychopathol*, *16*(2), 421-440. https://www.ncbi.nlm.nih.gov/pubmed/15487604
- Capaldi, D. M., & Rothbart, M. K. (1992). Development and validation of an early adolescent temperament measure. *Journal of Early Adolescence*, *12*(2), 153-173.
- Ellis, L. K., & Rothbart, M. K. (2001). Revision of the Early Adolescent Temperament Questionnaire. Poster presented at the 2001 Biennial Meeting of the Society for Research in Child Development, Minneapolis, Minnesota.
- **The questionnaire is available for research purposes upon request at https://research.bowdoin.edu/rothbart-temperament-questionnaires/ - see Parent-Report**

NOTE: Not all subscales were collected at Session 1 (ses-01). Gender neutral pronouns (and corresponding verb) were used at session 2 (ses-02), but not at session 1.

| QTAB Variable | Questionnaire Item Source | Comment |
|---|---|---|
| **pEATQ01** | Bowdoin website parent-report, Item 1 | |
| **pEATQ02** | Bowdoin website parent-report, Item 2 | **ses-02** “s/he” replaced with “they” |
| **pEATQ03** | Bowdoin website parent-report, Item 3 | |
| **pEATQ04** | Bowdoin website parent-report, Item 4 | |
| **pEATQ05** | Bowdoin website parent-report, Item 5 | |
| **pEATQ06** | Bowdoin website parent-report, Item 6 | **ses-02** “his/her” replaced with “their” |
| **pEATQ07** | Bowdoin website parent-report, Item 7 | **ses-02** “his/her” replaced with “their” |
| **pEATQ08** | Bowdoin website parent-report, Item 8 | **ses-02** “s/he” replaced with “they” |
| **pEATQ09** | Bowdoin website parent-report, Item 9 | |
| **pEATQ10** | Bowdoin website parent-report, Item 10 | |
| **pEATQ11** | Bowdoin website parent-report, Item 11 | |
| **pEATQ12** | Bowdoin website parent-report, Item 12 | |
| **pEATQ13** | Bowdoin website parent-report, Item 13 | |
| **pEATQ14** | Bowdoin website parent-report, Item 14 | **ses-02** “his/her” replaced with “their”; “s/he” replaced with “they” |
| **pEATQ15** | Bowdoin website parent-report, Item 15 | |
| **pEATQ16** | Bowdoin website parent-report, Item 16 | |
| **pEATQ17** | Bowdoin website parent-report, Item 17 | **ses-02** “s/he” replaced with “they” |
| **pEATQ18** | Bowdoin website parent-report, Item 18 | |
| **pEATQ19** | Bowdoin website parent-report, Item 19 | **ses-02** “s/he” replaced with “they” |
| **pEATQ20** | Bowdoin website parent-report, Item 20 | |
| **pEATQ21** | Bowdoin website parent-report, Item 21 | **ses-02** “him/her” replaced with “them” |
| **pEATQ22** | Bowdoin website parent-report, Item 22 | **ses-02** “s/he” replaced with “they” |
| **pEATQ23** | Bowdoin website parent-report, Item 23 | **ses-02** “s/he” replaced with “they”; “her/himself” replaced with “themself” |
| **pEATQ24** | Bowdoin website parent-report, Item 24 | **ses-02** “s/he” replaced with “they” |

| | | |
|---|---|---|
| **pEATQ25** | Bowdoin website parent-report, Item 25 | |
| **pEATQ26** | Bowdoin website parent-report, Item 26 | |
| **pEATQ27** | Bowdoin website parent-report, Item 27 | |
| **pEATQ28** | Bowdoin website parent-report, Item 28 | |
| **pEATQ29** | Bowdoin website parent-report, Item 29 | **ses-02** "s/he" replaced with "they" |
| **pEATQ30** | Bowdoin website parent-report, Item 30 | **ses-02** "s/he" replaced with "they" |
| **pEATQ31** | Bowdoin website parent-report, Item 31 | **ses-02** "him/her" replaced with "them"; "s/he" replaced with "they" |
| **pEATQ32** | Bowdoin website parent-report, Item 32 | |
| **pEATQ33** | Bowdoin website parent-report, Item 33 | |
| **pEATQ34** | Bowdoin website parent-report, Item 34 | |
| **pEATQ35** | Bowdoin website parent-report, Item 35 | |
| **pEATQ36** | Bowdoin website parent-report, Item 36 | **ses-02** "his/her" replaced with "their" |
| **pEATQ37** | Bowdoin website parent-report, Item 37 | |
| **pEATQ38** | Bowdoin website parent-report, Item 38 | |
| **pEATQ39** | Bowdoin website parent-report, Item 39 | **ses-02** "him/her" replaced with "them" |
| **pEATQ40** | Bowdoin website parent-report, Item 40 | |
| **pEATQ41** | Bowdoin website parent-report, Item 41 | |
| **pEATQ42** | Bowdoin website parent-report, Item 42 | |
| **pEATQ43** | Bowdoin website parent-report, Item 43 | |
| **pEATQ44** | Bowdoin website parent-report, Item 44 | |
| **pEATQ45** | Bowdoin website parent-report, Item 45 | **ses-02** "s/he" replaced with "they" |
| **pEATQ46** | Bowdoin website parent-report, Item 46 | |
| **pEATQ47** | Bowdoin website parent-report, Item 47 | **ses-02** "him/herself" replaced with "themself" |
| **pEATQ48** | Bowdoin website parent-report, Item 48 | **ses-02** "her/him" replaced with "them" |
| **pEATQ49** | Bowdoin website parent-report, Item 49 | |
| **pEATQ50** | Bowdoin website parent-report, Item 50 | |
| **pEATQ51** | Bowdoin website parent-report, Item 51 | |
| **pEATQ52** | Bowdoin website parent-report, Item 52 | **ses-02** "s/he" replaced with "they"; "her/himself" replaced with "themself" |
| **pEATQ53** | Bowdoin website parent-report, Item 53 | **ses-02** "s/he" replaced with "they" |
| **pEATQ54** | Bowdoin website parent-report, Item 54 | |
| **pEATQ55** | Bowdoin website parent-report, Item 55 | |
| **pEATQ56** | Bowdoin website parent-report, Item 56 | |
| **pEATQ57** | Bowdoin website parent-report, Item 57 | **ses-02** "him/her" replaced with "them" |
| **pEATQ58** | Bowdoin website parent-report, Item 58 | **ses-02** "s/he" replaced with "they"; "her/his" replaced with "their" |
| **pEATQ59** | Bowdoin website parent-report, Item 59 | **ses-02** "his/her" replaced with "their" |
| **pEATQ60** | Bowdoin website parent-report, Item 60 | |
| **pEATQ61** | Bowdoin website parent-report, Item 61 | |
| **pEATQ62** | Bowdoin website parent-report, Item 62 | |

**Social Support and Family Functioning**

### Multidimensional Scale of Perceived Social Support (MSPSS)

- Zimet, G. D., Dahlem, N. W., Zimet, S. G., & Farley, G. K. (1988). The Multidimensional Scale of Perceived Social Support. *Journal of Personality Assessment*, *52*(1), 30-41. https://doi.org/10.1207/s15327752jpa5201_2

| QTAB Variable | Scale Item Source | Comment |
|---|---|---|
| **MSPSS01** | Zimet et al 1988, Table 1, Item 1 | |
| **MSPSS02** | Zimet et al 1988, Table 1, Item 2 | |
| **MSPSS03** | Zimet et al 1988, Table 1, Item 3 | |
| **MSPSS04** | Zimet et al 1988, Table 1, Item 4 | |
| **MSPSS05** | Zimet et al 1988, Table 1, Item 5 | |
| **MSPSS06** | Zimet et al 1988, Table 1, Item 6 | |
| **MSPSS07** | Zimet et al 1988, Table 1, Item 7 | |
| **MSPSS08** | Zimet et al 1988, Table 1, Item 8 | |
| **MSPSS09** | Zimet et al 1988, Table 1, Item 9 | |
| **MSPSS10** | Zimet et al 1988, Table 1, Item 10 | |
| **MSPSS11** | Zimet et al 1988, Table 1, Item 11 | |
| **MSPSS12** | Zimet et al 1988, Table 1, Item 12 | |

### Alabama Parenting Questionnaire (APQ)

- Shelton, K. K., Frick, P. J., & Wootton, J. (1996). Assessment of parenting practices in families of elementary school-age children. *Journal of Clinical Child Psychology*, *25*(3), 317-329. https://doi.org/DOI 10.1207/s15374424jccp2503_8

NOTE: Not all subscales were collected at session 1 (only Poor Monitoring/Supervision) or session 2 (only Involvement, Positive Parenting, and Poor Monitoring/Supervision). Minor wording changes were made as questions are posed for more than one child (e.g. “child” is replaced with “children”; “his/her” is replaced with “their” etc.). Parent responds separately for each child.

| QTAB Variable | Questionnaire Item Source | Comment |
|---|---|---|
| **pAPQ01** | Shelton et al 1996, Table 2, Involvement, Item 1 | |
| **pAPQ02** | Shelton et al 1996, Table 2, Positive Parenting, Item 2 | “is doing a good job with something” inadvertently replaced with “do something well” |
| **pAPQ04** | Shelton et al 1996, Table 2, Involvement, Item 4 | |
| **pAPQ05** | Shelton et al 1996, Table 2, Positive Parenting, Item 5 | |
| **pAPQ06** | Shelton et al 1996, Table 2, Poor Monitoring, Item 6 | |
| **pAPQ07** | Shelton et al 1996, Table 2, Involvement, Item 7 | |
| **pAPQ09** | Shelton et al 1996, Table 2, Involvement, Item 9 | |
| **pAPQ10** | Shelton et al 1996, Table 2, Poor Monitoring, Item 10 | |
| **pAPQ11** | Shelton et al 1996, Table 2, Involvement, Item 11 | |
| **pAPQ13** | Shelton et al 1996, Table 2, Positive Parenting, Item 13 | |
| **pAPQ14** | Shelton et al 1996, Table 2, Involvement, Item 14 | |
| **pAPQ15** | Shelton et al 1996, Table 2, Involvement, Item 15 | “drive” replaced with “take” as some families use other means of transport |
| **pAPQ16** | Shelton et al 1996, Table 2, Positive Parenting, Item 16 | |
| **pAPQ17** | Shelton et al 1996, Table 2, Poor Monitoring, Item 17 | |
| **pAPQ18** | Shelton et al 1996, Table 2, Positive Parenting, Item 18 | |
| **pAPQ19** | Shelton et al 1996, Table 2, Poor Monitoring, Item 19 | |
| **pAPQ20** | Shelton et al 1996, Table 2, Involvement, Item 20 | |
| **pAPQ21** | Shelton et al 1996, Table 2, Poor Monitoring, Item 21 | |
| **pAPQ23** | Shelton et al 1996, Table 2, Involvement, Item 23 | |
| **pAPQ24** | Shelton et al 1996, Table 2, Poor Monitoring, Item 24 | |
| **pAPQ26** | Shelton et al 1996, Table 2, Involvement, Item 26 | |
| **pAPQ27** | Shelton et al 1996, Table 2, Positive Parenting, Item 27 | |
| **pAPQ28** | Shelton et al 1996, Table 2, Poor Monitoring, Item 28 | |
| **pAPQ29** | Shelton et al 1996, Table 2, Poor Monitoring, Item 29 | |
| **pAPQ30** | Shelton et al 1996, Table 2, Poor Monitoring, Item 30 | |
| **pAPQ32** | Shelton et al 1996, Table 2, Poor Monitoring, Item 32 | |

### McMaster Family Assessment Device (FAD)

- Epstein, N. B., Baldwin, L. M., & Bishop, D. S. (1983). THE McMASTER FAMILY ASSESSMENT DEVICE*. *Journal of Marital and Family Therapy*, *9*(2), 171-180. https://doi.org/10.1111/j.1752-0606.1983.tb01497.x
- Kabacoff, R. I., Miller, I. W., Bishop, D. S., Epstein, N. B., & Keitner, G. I. (1990). A psychometric study of the McMaster Family Assessment Device in psychiatric, medical, and nonclinical samples. *Journal of Family Psychology*, *3*(4), 431-439. https://doi.org/10.1037/h0080547

***For more about the 12-item General Functioning subscale see***

- **Boterhoven de Haan, K. L., Hafekost, J., Lawrence, D., Sawyer, M. G., & Zubrick, S. R. (2015). Reliability and validity of a short version of the general functioning subscale of the McMaster Family Assessment Device. *Fam Process*, *54*(1), 116-123. https://doi.org/10.1111/famp.12113**

**NOTE: the 6-time subscale used in Boterhoven de Haan et al 2015 is a match to QTAB items pFAD06, pFAD16, pFAD26, pFAD36, pFAD46, pFAD56.**

*These items were reverse scored (note that we were unable to obtain official identification of items to be reverse scored).

| QTAB Variable | Device Item Source | Comment |
|---|---|---|
| **pFAD01*** | Buckland Thesis, Appendix 3, Item 1<br>Epstein et al 1983, Table 1, General, 1st Item | |
| **pFAD02** | Buckland Thesis, Appendix 3, Item 2 | This item contributed to the “Problem Solving” Scale (personal communication with Sharon Buckland) |
| **pFAD03** | Buckland Thesis, Appendix 3, Item 3<br>Epstein et al 1983, Table 1, Communication, 1st Item | |
| **pFAD04*** | Buckland Thesis, Appendix 3, Item 4<br>Epstein et al 1983, Table 1, Roles, 1st Item | |
| **pFAD05*** | Buckland Thesis, Appendix 3, Item 5<br>Epstein et al 1983, Table 1, Affect Involvement, 1st Item | |
| **pFAD06** | Buckland Thesis, Appendix 3, Item 6<br>Epstein et al 1983, Table 1, General, 2nd Item | |
| **pFAD07*** | Buckland Thesis, Appendix 3, Item 7<br>Epstein et al 1983, Table 1, Behavior Control, 1st Item | |
| **pFAD08*** | Buckland Thesis, Appendix 3, Item 8 | This item contributed to the “Roles” Scale (personal communication with Sharon Buckland) |
| **pFAD09*** | Buckland Thesis, Appendix 3, Item 9<br>Epstein et al 1983, Table 1, Affect Resp, 1st Item | |
| **pFAD10** | Buckland Thesis, Appendix 3, Item 10<br>Epstein et al 1983, Table 1, Roles, 2nd Item | |
| **pFAD11*** | Buckland Thesis, Appendix 3, Item 11<br>Epstein et al 1983, Table 1, General, 3rd Item | |
| **pFAD12** | Buckland Thesis, Appendix 3, Item 12<br>Epstein et al 1983, Table 1, Problem Solving, 1st Item | |
| **pFAD13*** | Buckland Thesis, Appendix 3, Item 13<br>Epstein et al 1983, Table 1, Affect Involvement, 2nd Item | |
| **pFAD14*** | Buckland Thesis, Appendix 3, Item 14<br>Epstein et al 1983, Table 1, Communication, 2nd Item | |
| **pFAD15*** | Buckland Thesis, Appendix 3, Item 15<br>Epstein et al 1983, Table 1, Roles, 3rd Item | |
| **pFAD16** | Buckland Thesis, Appendix 3, Item 16<br>Epstein et al 1983, Table 1, General, 4th Item | |
| **pFAD17*** | Buckland Thesis, Appendix 3, Item 17<br>Epstein et al 1983, Table 1, Behavior Control, 2nd Item | |
| **pFAD18** | Buckland Thesis, Appendix 3, Item 18<br>Epstein et al 1983, Table 1, Communication, 3rd Item | |
| **pFAD19*** | Buckland Thesis, Appendix 3, Item 19<br>Epstein et al 1983, Table 1, Affect Resp, 2nd Item | |
| **pFAD20** | Buckland Thesis, Appendix 3, Item 20<br>Epstein et al 1983, Table 1, Behavior Control, 3rd Item | |
| **pFAD21*** | Buckland Thesis, Appendix 3, Item 21 | |

| | | |
|---|---|---|
| | Epstein et al 1983, Table 1, General, 5th Item | |
| **pFAD22*** | Buckland Thesis, Appendix 3, Item 22 | This item contributed to the "Communication" Scale (personal communication with Sharon Buckland) |
| **pFAD23*** | Buckland Thesis, Appendix 3, Item 23<br>Epstein et al 1983, Table 1, Roles, 4th Item | Wording as per Epstein et al 1983 |
| **pFAD24** | Buckland Thesis, Appendix 3, Item 24<br>Epstein et al 1983, Table 1, Problem Solving, 2nd Item | |
| **pFAD25*** | Buckland Thesis, Appendix 3, Item 25<br>Epstein et al 1983, Table 1, Affect Involvement, 3rd Item | |
| **pFAD26** | Buckland Thesis, Appendix 3, Item 26<br>Epstein et al 1983, Table 1, General, 6th Item | |
| **pFAD27*** | Buckland Thesis, Appendix 3, Item 27<br>Epstein et al 1983, Table 1, Behavior Control, 4th Item | |
| **pFAD28*** | Buckland Thesis, Appendix 3, Item 28<br>Epstein et al 1983, Table 1, Affect Resp, 3rd Item | |
| **pFAD29** | Buckland Thesis, Appendix 3, Item 29 | This item contributed to the "Communication" Scale (personal communication with Sharon Buckland) |
| **pFAD30** | Buckland Thesis, Appendix 3, Item 30 | This item contributed to the "Roles" Scale (personal communication with Sharon Buckland) |
| **pFAD31*** | Buckland Thesis, Appendix 3, Item 31<br>Epstein et al 1983, Table 1, General, 7th Item | |
| **pFAD32** | Buckland Thesis, Appendix 3, Item 32<br>Epstein et al 1983, Table 1, Behavior control, 5th Item | |
| **pFAD33*** | Buckland Thesis, Appendix 3, Item 33<br>Epstein et al 1983, Table 1, Affect Involvement, 4th Item | |
| **pFAD34*** | Buckland Thesis, Appendix 3, Item 34<br>Epstein et al 1983, Table 1, Roles, 5th Item | |
| **pFAD35*** | Buckland Thesis, Appendix 3, Item 35 | This item contributed to the "Communication" Scale (personal communication with Sharon Buckland) |
| **pFAD36** | Buckland Thesis, Appendix 3, Item 36<br>Epstein et al 1983, Table 1, General, 8th Item | |
| **pFAD37*** | Buckland Thesis, Appendix 3, Item 37<br>Epstein et al 1983, Table 1, Affect Involvement, 5th Item | |
| **pFAD38** | Buckland Thesis, Appendix 3, Item 38<br>Epstein et al 1983, Table 1, Problem Solving, 3rd Item | |
| **pFAD39*** | Buckland Thesis, Appendix 3, Item 39<br>Epstein et al 1983, Table 1, Affect Resp, 4th Item | |
| **pFAD40** | Buckland Thesis, Appendix 3, Item 40<br>Epstein et al 1983, Table 1, Roles, 6th Item | |
| **pFAD41*** | Buckland Thesis, Appendix 3, Item 41<br>Epstein et al 1983, Table 1, General, 9th Item | |
| **pFAD42*** | Buckland Thesis, Appendix 3, Item 42<br>Epstein et al 1983, Table 1, Affect Involvement, 6th Item | |
| **pFAD43** | Buckland Thesis, Appendix 3, Item 43<br>Epstein et al 1983, Table 1, Communication, 4th Item | Wording as per Epstein et al 1983 |
| **pFAD44*** | Buckland Thesis, Appendix 3, Item 44<br>Epstein et al 1983, Table 1, Behavior Control, 6th Item | |
| **pFAD45*** | Buckland Thesis, Appendix 3, Item 45<br>Epstein et al 1983, Table 1, Roles, 7th Item | |
| **pFAD46** | Buckland Thesis, Appendix 3, Item 46<br>Epstein et al 1983, Table 1, General, 10th Item | |
| **pFAD47*** | Buckland Thesis, Appendix 3, Item 47<br>Epstein et al 1983, Table 1, Behavior control, 7th Item | |
| **pFAD48*** | Buckland Thesis, Appendix 3, Item 48<br>Epstein et al 1983, Table 1, Behavior Control, 7th Item | |

| | | |
|---|---|---|
| **pFAD49** | Buckland Thesis, Appendix 3, Item 49<br>Epstein et al 1983, Table 1, Affect Resp, 5th Item | |
| **pFAD50** | Buckland Thesis, Appendix 3, Item 50<br>Epstein et al 1983, Table 1, Problem Solving, 4th Item | |
| **pFAD51*** | Buckland Thesis, Appendix 3, Item 51<br>Epstein et al 1983, Table 1, General, 11th Item | |
| **pFAD52*** | Buckland Thesis, Appendix 3, Item 52<br>Epstein et al 1983, Table 1, Communication, 5th Item | |
| **pFAD53*** | Buckland Thesis, Appendix 3, Item 53<br>Epstein et al 1983, Table 1, Roles, 8th Item | |
| **pFAD54*** | Buckland Thesis, Appendix 3, Item 54<br>Epstein et al 1983, Table 1, Affect Involvement, 7th Item | |
| **pFAD55** | Buckland Thesis, Appendix 3, Item 55<br>Epstein et al 1983, Table 1, Behavior Control, 9th Item | |
| **pFAD56** | Buckland Thesis, Appendix 3, Item 56<br>Epstein et al 1983, Table 1, General, 12th Item | |
| **pFAD57** | Buckland Thesis, Appendix 3, Item 57<br>Epstein et al 1983, Table 1, Affect Resp, 6th Item | |
| **pFAD58*** | Buckland Thesis, Appendix 3, Item 58 | This item contributed to the "Roles" Scale (personal communication with Sharon Buckland) |
| **pFAD59** | Buckland Thesis, Appendix 3, Item 59<br>Epstein et al 1983, Table 1, Communication, 6th Item | |
| **pFAD60** | Buckland Thesis, Appendix 3, Item 60<br>Epstein et al 1983, Table 1, Problem Solving, 5th Item | |

**Stress**

### Daily Life Stressors Scale (DLSS)

- Kearney, C. A., Drabman, R. S., & Beasley, J. F. (1993). The trials of childhood: the development, reliability, and validity of the Daily Life Stressors Scale. *J Child Fam Stud*, *2*(4), 371-388. https://doi.org/10.1007/BF01321232

| QTAB Variable | Scale Item Source | Comment |
|---|---|---|
| **DLSS01** | Kearney et al 1993, Table 1, Item 1 | |
| **DLSS02** | Kearney et al 1993, Table 1, Item 2 | |
| **DLSS03** | Kearney et al 1993, Table 1, Item 3 | |
| **DLSS04** | Kearney et al 1993, Table 1, Item 4 | |
| **DLSS05** | Kearney et al 1993, Table 1, Item 5 | |
| **DLSS06** | Kearney et al 1993, Table 1, Item 6 | |
| **DLSS07** | Kearney et al 1993, Table 1, Item 7 | |
| **DLSS08** | Kearney et al 1993, Table 1, Item 8 | |
| **DLSS09** | Kearney et al 1993, Table 1, Item 9 | |
| **DLSS10** | Kearney et al 1993, Table 1, Item 10 | |
| **DLSS11** | Kearney et al 1993, Table 1, Item 11 | |
| **DLSS12** | Kearney et al 1993, Table 1, Item 12 | |
| **DLSS13** | Kearney et al 1993, Table 1, Item 13 | |
| **DLSS14** | Kearney et al 1993, Table 1, Item 14 | |
| **DLSS15** | Kearney et al 1993, Table 1, Item 15 | |
| **DLSS16** | Kearney et al 1993, Table 1, Item 16 | |
| **DLSS17** | Kearney et al 1993, Table 1, Item 17 | |
| **DLSS18** | Kearney et al 1993, Table 1, Item 18 | |
| **DLSS19** | Kearney et al 1993, Table 1, Item 19 | |
| **DLSS20** | Kearney et al 1993, Table 1, Item 20 | |
| **DLSS21** | Kearney et al 1993, Table 1, Item 21 | |
| **DLSS22** | Kearney et al 1993, Table 1, Item 22 | |
| **DLSS23** | Kearney et al 1993, Table 1, Item 23 | |
| **DLSS24** | Kearney et al 1993, Table 1, Item 24 | |
| **DLSS25** | Kearney et al 1993, Table 1, Item 25 | |
| **DLSS26** | Kearney et al 1993, Table 1, Item 26 | |
| **DLSS27** | Kearney et al 1993, Table 1, Item 27 | |
| **DLSS28** | Kearney et al 1993, Table 1, Item 28 | |
| **DLSS29** | Kearney et al 1993, Table 1, Item 29 | |
| **DLSS30** | Kearney et al 1993, Table 1, Item 30 | |

### Gatehouse Bullying Scale (GBS)

- Bond, L., Wolfe, S., Tollit, M., Butler, H., & Patton, G. (2007). A comparison of the Gatehouse Bullying Scale and the peer relations questionnaire for students in secondary school. *J Sch Health*, *77*(2), 75-79. https://doi.org/10.1111/j.1746-1561.2007.00170.x

**Data were scored as described in:**

- Thomas, H. J., Chan, G. C., Scott, J. G., Connor, J. P., Kelly, A. B., & Williams, J. (2016). Association of different forms of bullying victimisation with adolescents' psychological distress and reduced emotional wellbeing. *Aust N Z J Psychiatry*, *50*(4), 371-379. https://doi.org/10.1177/0004867415600076

| QTAB Variable | Scale Item Source | Comment |
|---|---|---|
| **GBS01a** | Bond et al 2007, Table 1, GBS, Item 1a | |
| **GBS01b** | Bond et al 2007, Table 1, GBS, Item 1b | |
| **BGS01c** | Bond et al 2007, Table 1, GBS, Item 1c | |
| **GBS02a** | Bond et al 2007, Table 1, GBS, Item 2a | |
| **GBS02b** | Bond et al 2007, Table 1, GBS, Item 2b | |
| **BGS02c** | Bond et al 2007, Table 1, GBS, Item 2c | |
| **GBS03a** | Bond et al 2007, Table 1, GBS, Item 3a | |
| **GBS03b** | Bond et al 2007, Table 1, GBS, Item 3b | |
| **BGS03c** | Bond et al 2007, Table 1, GBS, Item 3c | |
| **GBS04a** | Bond et al 2007, Table 1, GBS, Item 4a | |
| **GBS04b** | Bond et al 2007, Table 1, GBS, Item 4b | |
| **BGS04c** | Bond et al 2007, Table 1, GBS, Item 4c | |

### Childhood Life Events Questionnaire (CLEQ)

- Upthegrove, R., Chard, C., Jones, L., Gordon-Smith, K., Forty, L., Jones, I., & Craddock, N. (2015). Adverse childhood events and psychosis in bipolar affective disorder. *Br J Psychiatry*, *206*(3), 191-197. https://doi.org/10.1192/bjp.bp.114.152611

Note: The CLEQ was used in questionnaire format and was answered by a parent for each adolescent twin. Each item was prefaced with “Have your children experienced: …”.

| QTAB Variable | Questionnaire Item Source | Comment |
|---|---|---|
| **pCLEQ01** | Upthegrove et al 2015, Data Supplement, Item 1 | |
| **pCLEQ01a** | If yes to Item 1, at what age? | |
| **pCLEQ02** | Upthegrove et al 2015, Data Supplement, Item 2 | |
| **pCLEQ02a** | If yes to Item 2, at what age? | |
| **pCLEQ03** | Upthegrove et al 2015, Data Supplement, Item 3 | |
| **pCLEQ03a** | If yes to Item 3, at what age? | |
| **pCLEQ04** | Upthegrove et al 2015, Data Supplement, Item 4 | |
| **pCLEQ04a** | If yes to Item 4, at what age? | |
| **pCLEQ05** | Upthegrove et al 2015, Data Supplement, Item 5 | |
| **pCLEQ05a** | If yes to Item 5, at what age? | |
| **pCLEQ06** | Upthegrove et al 2015, Data Supplement, Item 6 | |
| **pCLEQ06a** | If yes to Item 6, at what age? | |
| **pCLEQ07** | Upthegrove et al 2015, Data Supplement, Item 7 | |
| **pCLEQ07a** | If yes to Item 7, at what age? | |
| **pCLEQ08** | Upthegrove et al 2015, Data Supplement, Item 8 | |
| **pCLEQ08a** | If yes to Item 8, at what age? | |
| **pCLEQ09** | Upthegrove et al 2015, Data Supplement, Item 9 | |
| **pCLEQ09a** | If yes to Item 9, at what age? | |
| **pCLEQ10** | Upthegrove et al 2015, Data Supplement, Item 10 | |
| **pCLEQ10a** | If yes to Item 10, at what age? | |
| | Upthegrove et al 2015, Data Supplement, Item 11 | This item was not included. |
| **pCLEQ12** | Upthegrove et al 2015, Data Supplement, Item 12 | |
| **pCLEQ12a** | If yes to Item 12, at what age? | |
| | Upthegrove et al 2015, Data Supplement, Item 13 | This item not available. |

### Prenatal Stress Exposure Scale (PSES)

**Adapted from interview reported in**

- Favaro, A., Tenconi, E., Degortes, D., Manara, R., & Santonastaso, P. (2015). Neural correlates of prenatal stress in young women. *Psychol Med*, *45*(12), 2533-2543. https://doi.org/10.1017/S003329171500046X

| QTAB Variable | Scale Items<br>(Were any of the following traumatic or stressful events experienced **in the 12 months before the twin’s conception, during the pregnancy, or in the first months after the birth of the twins**?) | Comment |
|---|---|---|
| **pPSES01a** | Did this occur: Accidents (including those of a close friend or relative) | |
| **pPSES01b** | If yes, when did the event occur (can select multiple)? | • In 12 months before conception<br>• During the pregnancy |

| | | |
|---|---|---|
| | | • After delivery |
| **pPSES01c** | If yes, what level of stress did the (most stressful) event cause? | • No significant stress<br>• Some stress<br>• Moderate stress<br>• Substantial stress<br>• Extreme stress |
| **pPSES02a** | Did this occur: Death of a friend or close relative | |
| **pPSES02b** | If yes, when did the event occur (can select multiple)? | |
| **pPSES02c** | If yes, what level of stress did the (most stressful) event cause? | |
| **pPSES03a** | Did this occur: Health problems (unrelated to the pregnancy) | |
| **pPSES03b** | If yes, when did the event occur (can select multiple)? | |
| **pPSES03c** | If yes, what level of stress did the (most stressful) event cause? | |
| **pPSES04a** | Did this occur: Severe health problem of a close friend or relative, or need to give assistance to a sick friend or relative | |
| **pPSES04b** | If yes, when did the event occur (can select multiple)? | |
| **pPSES04c** | If yes, what level of stress did the (most stressful) event cause? | |
| **pPSES05a** | Did this occur: Natural disasters (e.g. flooding) | |
| **pPSES05b** | If yes, when did the event occur (can select multiple)? | |
| **pPSES05c** | If yes, what level of stress did the (most stressful) event cause? | |
| **pPSES06a** | Did this occur: Severe interpersonal conflicts with a partner or close relative | |
| **pPSES06b** | If yes, when did the event occur (can select multiple)? | |
| **pPSES06c** | If yes, what level of stress did the (most stressful) event cause? | |
| **pPSES07a** | Did this occur: Separation from a partner | |
| **pPSES07b** | If yes, when did the event occur (can select multiple)? | |
| **pPSES07c** | If yes, what level of stress did the (most stressful) event cause? | |
| **pPSES08a** | Did this occur: Severe legal or economic problems | |
| **pPSES08b** | If yes, when did the event occur (can select multiple)? | |
| **pPSES08c** | If yes, what level of stress did the (most stressful) event cause? | |
| **pPSES09a** | Did this occur: Relocation (i.e. moving house) | |
| **pPSES09b** | If yes, when did the event occur (can select multiple)? | |
| **pPSES09c** | If yes, what level of stress did the (most stressful) event cause? | |
| **pPSES10a** | Did this occur: Personal violence | |
| **pPSES10b** | If yes, when did the event occur (can select multiple)? | |
| **pPSES10c** | If yes, what level of stress did the (most stressful) event cause? | |
| **pPSES11a** | Did this occur: Sexual abuse or maltreatment | |
| **pPSES11b** | If yes, when did the event occur (can select multiple)? | |
| **pPSES11c** | If yes, what level of stress did the (most stressful) event cause? | |
| **pPSES12a** | Did this occur: Miscarriage or abortion (occurring in the 12 months before the twin's conception) | |
| **pPSES12c** | If yes, what level of stress did the (most stressful) event cause? | |
| **pPSES13a** | Did this occur: Other offspring death | |
| **pPSES13b** | If yes, when did the event occur (can select multiple)? | |
| **pPSES13c** | If yes, what level of stress did the (most stressful) event cause? | |
| **pPSES14a** | Did this occur: Other traumatic or stressful event | |
| **pPSES14b** | If yes, when did the event occur (can select multiple)? | |

| | |
|---|---|
| **pPSES14c** | If yes, what level of stress did the (most stressful) event cause? |

### Parental Stress Scale (PaSS)

- Berry, J. O., & Jones, W. H. (1995). The Parental Stress Scale: initial psychometric evidence. *Journal of Social and Personal Relationships*, *12*(3), 463-472. https://doi.org/10.1177/0265407595123009

| QTAB Variable | Scale Item Source | Comment |
|---|---|---|
| **pPaSS01** | Berry et al 1995, Table 1, Item 1 | |
| **pPaSS02** | Berry et al 1995, Table 1, Item 2 | |
| **pPaSS03** | Berry et al 1995, Table 1, Item 3 | |
| **pPaSS04** | Berry et al 1995, Table 1, Item 4 | |
| **pPaSS05** | Berry et al 1995, Table 1, Item 5 | |
| **pPaSS06** | Berry et al 1995, Table 1, Item 6 | |
| **pPaSS07** | Berry et al 1995, Table 1, Item 7 | |
| **pPaSS08** | Berry et al 1995, Table 1, Item 8 | |
| **pPaSS09** | Berry et al 1995, Table 1, Item 9 | |
| **pPaSS10** | Berry et al 1995, Table 1, Item 10 | |
| **pPaSS11** | Berry et al 1995, Table 1, Item 11 | |
| **pPaSS12** | Berry et al 1995, Table 1, Item 12 | |
| **pPaSS13** | Berry et al 1995, Table 1, Item 13 | |
| **pPaSS14** | Berry et al 1995, Table 1, Item 14 | |
| **pPaSS15** | Berry et al 1995, Table 1, Item 15 | |
| **pPaSS16** | Berry et al 1995, Table 1, Item 16 | |
| **pPaSS17** | Berry et al 1995, Table 1, Item 17 | |
| **pPaSS18** | Berry et al 1995, Table 1, Item 18 | |

### List of Threatening Experiences (LTE)

- Motrico, E., Moreno-Kustner, B., de Dios Luna, J., Torres-Gonzalez, F., King, M., Nazareth, I., Monton-Franco, C., Gilde Gomez-Barragan, M. J., Sanchez-Celaya, M., Diaz-Barreiros, M. A., Vicens, C., Moreno-Peral, P., & Bellon, J. A. (2013). Psychometric properties of the List of Threatening Experiences--LTE and its association with psychosocial factors and mental disorders according to different scoring methods. *J Affect Disord*, *150*(3), 931-940. https://doi.org/10.1016/j.jad.2013.05.017

NOTE: all items are prefaced with “during the last 12 months, have you experienced…”

| QTAB Variable | Scale Item Source | Comment |
|---|---|---|
| **pLTE01** | Motrico et al 2013, Table 2, Item 1 | |
| **pLTE02** | Motrico et al 2013, Table 2, Item 2 | |
| **pLTE03** | Motrico et al 2013, Table 2, Item 3 | |
| **pLTE04** | Motrico et al 2013, Table 2, Item 4 | |
| **pLTE05** | Motrico et al 2013, Table 2, Item 5 | |
| **pLTE06** | Motrico et al 2013, Table 2, Item 6 | |
| **pLTE07** | Motrico et al 2013, Table 2, Item 7 | |
| **pLTE08** | Motrico et al 2013, Table 2, Item 8 | |
| **pLTE09** | Motrico et al 2013, Table 2, Item 9 | |
| **pLTE10** | Motrico et al 2013, Table 2, Item 10 | |
| **pLTE11** | Motrico et al 2013, Table 2, Item 11 | |
| **pLTE12** | Motrico et al 2013, Table 2, Item 12 | |

**Sleep and Physical Health**

### Pediatric Daytime Sleepiness Scale (PDSS)

- Drake, C., Nickel, C., Burduvali, E., Roth, T., Jefferson, C., & Pietro, B. (2003). The pediatric daytime sleepiness scale (PDSS): sleep habits and school outcomes in middle-school children. *Sleep*, *26*(4), 455-458. https://www.ncbi.nlm.nih.gov/pubmed/12841372

| QTAB Variable | Scale Item Source | Comment |
|---|---|---|
| **PDSS01** | Drake et al 2003, Appendix, Item 1 | |
| **PDSS02** | Drake et al 2003, Appendix, Item 2 | |
| **PDSS03** | Drake et al 2003, Appendix, Item 3 | |
| **PDSS04** | Drake et al 2003, Appendix, Item 4 | |
| **PDSS05** | Drake et al 2003, Appendix, Item 5 | |
| **PDSS06** | Drake et al 2003, Appendix, Item 6 | |
| **PDSS07** | Drake et al 2003, Appendix, Item 7 | |
| **PDSS08** | Drake et al 2003, Appendix, Item 8 | |

### Sleep Disturbances Scale for Children (SDSC)

- Bruni, O., Ottaviano, S., Guidetti, V., Romoli, M., Innocenzi, M., Cortesi, F., & Giannotti, F. (1996). The Sleep Disturbance Scale for Children (SDSC). Construction and validation of an instrument to evaluate sleep disturbances in childhood and adolescence. *J Sleep Res*, *5*(4), 251-261. https://doi.org/10.1111/j.1365-2869.1996.00251.x

NOTE: "The child" has been replaced with "your child" throughout. Gender neutral pronouns (and corresponding verb) were used.

| QTAB Variable | Scale Item Source | Comment |
|---|---|---|
| **pSDSC01** | Bruni et al 1996, Appendix A, Item 1 | |
| **pSDSC02** | Bruni et al 1996, Appendix A, Item 2 | |
| **pSDSC03** | Bruni et al 1996, Appendix A, Item 3 | |
| **pSDSC04** | Bruni et al 1996, Appendix A, Item 4 | |
| **pSDSC05** | Bruni et al 1996, Appendix A, Item 5 | |
| **pSDSC06** | Bruni et al 1996, Appendix A, Item 6 | |
| **pSDSC07** | Bruni et al 1996, Appendix A, Item 7 | |
| **pSDSC08** | Bruni et al 1996, Appendix A, Item 8 | |
| **pSDSC09** | Bruni et al 1996, Appendix A, Item 9 | |
| **pSDSC10** | Bruni et al 1996, Appendix A, Item 10 | |
| **pSDSC11** | Bruni et al 1996, Appendix A, Item 11 | |
| **pSDSC12** | Bruni et al 1996, Appendix A, Item 12 | |
| **pSDSC13** | Bruni et al 1996, Appendix A, Item 13 | |
| **pSDSC14** | Bruni et al 1996, Appendix A, Item 14 | |
| **pSDSC15** | Bruni et al 1996, Appendix A, Item 15 | |
| **pSDSC16** | Bruni et al 1996, Appendix A, Item 16 | |
| **pSDSC17** | Bruni et al 1996, Appendix A, Item 17 | |
| **pSDSC18** | Bruni et al 1996, Appendix A, Item 18 | "his/her" replaced with "their" |
| **pSDSC19** | Bruni et al 1996, Appendix A, Item 19 | |
| **pSDSC20** | Bruni et al 1996, Appendix A, Item 20 | "him/her" replaced with "them" |
| **pSDSC21** | Bruni et al 1996, Appendix A, Item 21 | "he/she" replaced with "they" |
| **pSDSC22** | Bruni et al 1996, Appendix A, Item 22 | |
| **pSDSC23** | Bruni et al 1996, Appendix A, Item 23 | |
| **pSDSC24** | Bruni et al 1996, Appendix A, Item 24 | |
| **pSDSC25** | Bruni et al 1996, Appendix A, Item 25 | "somnolence" was replaced with "sleepiness" |
| **pSDSC26** | Bruni et al 1996, Appendix A, Item 26 | |

### Sleep behaviours across early childhood

**Adapted from Generation R sleep items reported in:**

- Kocevska, D., Muetzel, R. L., Luik, A. I., Luijk, M. P., Jaddoe, V. W., Verhulst, F. C., White, T., & Tiemeier, H. (2017). The Developmental Course of Sleep Disturbances Across Childhood Relates to Brain Morphology at Age 7: The Generation R Study. *Sleep*, *40*(1). https://doi.org/10.1093/sleep/zsw022

Generation R items: Has trouble getting to sleep (QTAB item1); Sleeps less than most kids during day and/or night (QTAB item 2); Wakes up often at night (QTAB item 3); Doesn't want to sleep alone (QTAB item 4); Resists going to bed at night (QTAB item 5); Parental presence required at bedtime (QTAB item 6)

| QTAB Variable | Item | Comment |
|---|---|---|
| **pGenR_2mth** | Sleep behaviour sum score at age 2 months | Sum of variables<br>pGenR10 (Item 1 at age 2 months)<br>pGenR20 (Item 2 at age 2 months)<br>pGenR30 (Item 3 at age 2 months)<br>pGenR40 (Item 4 at age 2 months) |
| **pGenR_18mth** | Sleep behaviour sum score at age 18 months | Sum of variables<br>pGenR11 (Item 1 at age 18 months)<br>pGenR21 (Item 2 at age 18 months)<br>pGenR31 (Item 3 at age 18 months)<br>pGenR41 (Item 4 at age 18 months)<br>pGenR51 (Item 5 at age 18 months)<br>pGenR61 (Item 6 at age 18 months) |
| **pGenR_2yrs** | Sleep behaviour sum score at age 2 years | Sum of variables<br>pGenR12 (Item 1 at age 2 years)<br>pGenR22 (Item 2 at age 2 years)<br>pGenR32 (Item 3 at age 2 years)<br>pGenR42 (Item 4 at age 2 years) |

| | | |
|---|---|---|
| | | pGenR52 (Item 5 at age 2 years)<br>pGenR62 (Item 6 at age 2 years) |
| **pGenR_3yrs** | Sleep behaviour sum score at age 3 years | Sum of variables<br>pGenR13 (Item 1 at age 3 years)<br>pGenR23 (Item 2 at age 3 years)<br>pGenR33 (Item 3 at age 3 years)<br>pGenR43 (Item 4 at age 3 years)<br>pGenR53 (Item 5 at age 3 years)<br>pGenR63 (Item 6 at age 3 years) |
| **pGenR_6yrs** | Sleep behaviour sum score at age 6 years | Sum of variables<br>pGenR16 (Item 1 at age 6 years)<br>pGenR26 (Item 2 at age 6 years)<br>pGenR36 (Item 3 at age 6 years)<br>pGenR46 (Item 4 at age 6 years)<br>pGenR56 (Item 5 at age 6 years)<br>pGenR66 (Item 6 at age 6 years) |
| **pGenR_9yrs** | Sleep behaviour sum score at age 9 years | Sum of variables<br>pGenR19 (Item 1 at age 9 years)<br>pGenR29 (Item 2 at age 9 years)<br>pGenR39 (Item 3 at age 9 years)<br>pGenR49 (Item 4 at age 9 years)<br>pGenR59 (Item 5 at age 9 years)<br>pGenR69 (Item 6 at age 9 years) |

**COVID-19 Pandemic Specific Assessments (subsample only)**

### Active and Passive Social Media Use (APSMU)

- Thorisdottir, I. E., Sigurvinsdottir, R., Asgeirsdottir, B. B., Allegrante, J. P., & Sigfusdottir, I. D. (2019). Active and Passive Social Media Use and Symptoms of Anxiety and Depressed Mood Among Icelandic Adolescents. *Cyberpsychol Behav Soc Netw*, *22*(8), 535-542. https://doi.org/10.1089/cyber.2019.0079
- Frison, E., & Eggermont, S. (2015). Toward an Integrated and Differential Approach to the Relationships Between Loneliness, Different Types of Facebook Use, and Adolescents' Depressed Mood. *Communication Research*, *47*(5), 701-728. https://doi.org/10.1177/0093650215617506

| QTAB Variable | Item | Comment |
|---|---|---|
| **APSMU01** | Thorisdottir et al 2019, Table 1, 1st Item | |
| **APSMU02** | Thorisdottir et al 2019, Table 1, 2nd Item | |
| **APSMU03** | Thorisdottir et al 2019, Table 1, 3rd Item | |
| **APSMU04** | Thorisdottir et al 2019, Table 1, 4th Item | |
| **APSMU05** | Thorisdottir et al 2019, Table 1, 5th Item | |
| **APSMU06** | Thorisdottir et al 2019, Table 1, 6th Item | |

### UCLA Brief COVID-19 Screen for Child/Adolescent PTSD

- UCLA Brief COVID-19 Screen for Child/Adolescent PTSD. (2020). The Regents of the University of California.
- **https://istss.org/getattachment/Clinical-Resources/Assessing-Trauma/UCLA-Posttraumatic-Stress-Disorder-Reaction-Index/UCLA-Brief-COVID-19-Screening-Form-English-4-13-20.pdf** (accessed 24th November, 2022)

| QTAB Variable | Item | Comment |
|---|---|---|
| **PTSD01** | Online pdf, Page 2, Item 1 | |
| **PTSD02** | Online pdf, Page 2, Item 2 | |
| **PTSD03** | Online pdf, Page 2, Item 3 | |
| **PTSD04** | Online pdf, Page 2, Item 4 | |
| **PTSD05** | Online pdf, Page 2, Item 5 | |
| **PTSD06** | Online pdf, Page 2, Item 6 | |
| **PTSD07** | Online pdf, Page 2, Item 7 | |
| **PTSD08** | Online pdf, Page 2, Item 8 | |
| **PTSD09** | Online pdf, Page 2, Item 9 | |
| **PTSD010** | Online pdf, Page 2, Item 10 | |
| **PTSD011** | Online pdf, Page 2, Item 11 | |

### Perceived Stress Scale (PSS)

- Cohen, S., Kamarck, T., & Mermelstein, R. (1983). A global measure of perceived stress. *J Health Soc Behav*, *24*(4), 385-396. https://www.ncbi.nlm.nih.gov/pubmed/6668417
- Lee, B., & Jeong, H. I. (2019). Construct validity of the perceived stress scale (PSS-10) in a sample of early childhood teacher candidates. *Psychiatry and Clinical Psychopharmacology*, *29*(1), 76-82. https://doi.org/10.1080/24750573.2019.1565693

Note: the 10-item version of the PSS was used.

| QTAB Variable | Scale Item | Comment |
|---|---|---|
| **pPSS01** | Cohen et al 1983, Appendix A, Item 1<br>Lee et al 2019, Table 1, Item 1 | |
| **pPSS02** | Cohen et al 1983, Appendix A, Item 2 | |

| | | |
|---|---|---|
| | Lee et al 2019, Table 1, Item 2 | |
| **pPSS03** | Cohen et al 1983, Appendix A, Item 3<br>Lee et al 2019, Table 1, Item 3 | |
| **pPSS04** | Cohen et al 1983, Appendix A, Item 4<br>Lee et al 2019, Table 1, Item 4 | Lee et al 2019 version used |
| **pPSS05** | Cohen et al 1983, Appendix A, Item 7<br>Lee et al 2019, Table 1, Item 5 | Lee et al 2019 version used |
| **pPSS06** | Cohen et al 1983, Appendix A, Item 8<br>Lee et al 2019, Table 1, Item 6 | |
| **pPSS07** | Cohen et al 1983, Appendix A, Item 9<br>Lee et al 2019, Table 1, Item 7 | |
| **pPSS08** | Cohen et al 1983, Appendix A, Item 10<br>Lee et al 2019, Table 1, Item 8 | |
| **pPSS09** | Cohen et al 1983, Appendix A, Item 11<br>Lee et al 2019, Table 1, Item 9 | Lee et al 2019 version used |
| **pPSS10** | Cohen et al 1983, Appendix A, Item 14<br>Lee et al 2019, Table 1, Item 10 | |

### Brief Resilience Scale (BRS)

- Smith, B. W., Dalen, J., Wiggins, K., Tooley, E., Christopher, P., & Bernard, J. (2008). The brief resilience scale: assessing the ability to bounce back. *Int J Behav Med*, *15*(3), 194-200. https://doi.org/10.1080/10705500802222972

| QTAB Variable | Scale Item | Comment |
|---|---|---|
| **BRS01 (self-report)**<br>**pBRS01 (parent-report)** | Smith et al 2008, Table 1, Item 1 | |
| **BRS02**<br>**pBRS02** | Smith et al 2008, Table 1, Item 2 | |
| **BRS03**<br>**pBRS03** | Smith et al 2008, Table 1, Item 3 | |
| **BRS04**<br>**pBRS04** | Smith et al 2008, Table 1, Item 4 | |
| **BRS05**<br>**pBRS05** | Smith et al 2008, Table 1, Item 5 | |
| **BRS06**<br>**pBRS06** | Smith et al 2008, Table 1, Item 6 | |
